# MIMOSA: Guiding De Novo Antibody Design with Natural Interaction Fingerprints

**DOI:** 10.64898/2026.09.24.753155

**Authors:** Brennan Abanades, Andrea Roncoli, Maniraj Bhagawati, Pan Kessel, Sabine Imhof-Jung, Dominique Doerr, Jonas Schilz, Stefania Vasilaki, Franziska Seeger, Alexandre M.J.J. Bonvin, Richard Bonneau, Vladimir Gligorijevic, Anna Vangone

**Affiliations:** Prescient Design - AI for Drug Discovery, Computational Science Center of Excellence, Roche; Large Molecule Research, Roche Pharma Research and Early Development, Roche Innovation Center Munich, Penzberg, Germany; Utrecht University, Faculty of Science - Chemistry, Bijvoet Centre, the Netherlands

## Abstract

De novo antibody design promises to transform therapeutic antibody and nanobody discovery. Yet, success remains inconsistent across targets, with generative models often yielding no experimentally validated binders. Here we introduce MIMOSA (MIMic-Oriented Structural Antibody generation), a model-agnostic, inference-time framework that constrains diffusion-based antibody design pipelines to reproduce the interaction geometry and chemistry of a target’s known cognate binder interface. Rather than searching for a productive interface from scratch, MIMOSA constrains generation around residue identities and geometries already known to support binding, while allowing the generative model to complete the surrounding antibody interface. We test MIMOSA on different interface topologies: contiguous motifs that fit within a single CDR loop and spatially dispersed hotspots distributed across the paratope. Applied without retraining to two architecturally distinct generators (RFAntibody and BoltzGen), this framework delivers experimentally validated binders–achieving sub-micromolar to single-digit nanomolar affinities–at 20–26% hit rates on three targets on which unconstrained methods produce zero or near-zero binders: KEAP1, uPA, and IL-8. By turning any target with a structurally characterised cognate binder into an accessible de novo design problem, this framework broadens the practical reach of generative antibody design to targets where unconstrained approaches currently fail.

## 1 Introduction

With over 200 approved abtibody-based drugs globally and nearly 1,400 candidates in clinical evaluation (Crescioli et al.x, 2026), antibodies have become a dominant class of therapeutics. Their high target specificity, favourable pharmacokinetics, and ability to engage surfaces inaccessible to small molecules underpin a growing presence across oncology, immunology, and infectious diseases (Carter and Lazar, 2018). However, discovering antibodies that bind a defined epitope remains slow and uncertain. Conventional campaigns relying on animal immunization, phage or yeast display (Valldorf et al., 2022), and iterative affinity maturation can be extensive and time consuming, often requiring multiple cycles of optimization (Laustsen et al., 2021), and even then offer no guarantee that the recovered binders will engage the therapeutically relevant site. Epitope specificity is particularly difficult to engineer, often requiring extensive counter-selection or structural characterization to confirm the intended mechanism of action.

De novo antibody design can dramatically speed up large molecule drug discovery, with both diffusion-based and hallucination-based approaches producing antibodies and nanobodies computationally. Several systems are demonstrating experimentally validated binders: RFAntibody achieved the first structurally confirmed de novo VHHs targeting user-specified epitopes (Bennett et al., 2024), BoltzGen recovered nanomolar binders for the majority of novel targets from fewer than 15 tested designs per target (Stark et al., 2025), and Germinal and mBER have further expanded the accessible target space with open-source pipelines validated at scale (Mille-Fragoso et al., 2025; Swanson et al., 2025). Despite this progress, consistent hit rates from manageable numbers of designs remain challenging. Generative models can propose large numbers of structurally plausible binders, but only a small fraction achieve the precise residue identities, geometries, and interaction networks required for productive binding. The search space of possible designs is vast, and reliably identifying true binders within it has become a central bottleneck. Strategies that focus generation toward binding-relevant solutions, rather than sampling the full space, are therefore critical to improving experimental success.

Natural ligand-receptor interactions offer a source of guidance for both instantiation and scoring of de novo design. When a cognate ligand is known, its hotspot residues define a proven, transferable interaction fingerprint: the specific residue identities, positions, and geometries that drive productive binding. An early realisation of this idea was hotspot grafting: Liu et al. searched a library of natural antibody scaffolds for CDR positions whose geometry could host the fixed Nrf2 ETGE hotspot triad, yielding a Kelch-like ECH-associated protein 1 (KEAP1) binder (Liu et al., 2017). Immune repertoires can also converge on mimic-like solutions: the alpaca-derived nanobody Nb4 binds urokinase plasminogen activator (uPA) via a substrate-like CDR-H3 (Kromann-Hansen et al., 2016), and we recently showed that rabbit antibodies against interleukin-18 receptor alpha (IL-18RA) extend this mimicry across multiple CDR loops (Abanades et al., 2026). Such repertoire-driven solutions arrive through affinity maturation at the same fingerprints Liu et al. exploited by design. Here, we extend this principle to generative de novo design: rather than transplanting motifs into existing scaffolds or mining repertoires for pre-existing mimics, we guide generative models to build antibody backbones whose CDRs can naturally host the motif.

We call this framework MIMOSA: MIMic-Oriented Structural Antibody generation. MIMOSA acts at two stages of the de novo design pipeline. During diffusion-based backbone structure generation, a differentiable geometric potential, which we term the “mimic potential”, steers CDR coordinates toward interaction hotspots from a known target–partner complex (Fig. 1a,b). During inverse folding, the chemical identities of successfully placed hotspot residues are preserved while the remaining CDR sequence is designed. Unlike conventional motif scaffolding (Azoitei et al., 2011; Correia et al., 2014; Watson et al., 2023; Lin et al., 2026), the mimic potential does not require a pre-determined mapping between motif residues and output sequence positions. Instead, it dynamically assigns motif residues to geometrically compatible CDR positions during denoising, allowing the model to discover where within the CDR loops the motif naturally fits. The mimic potential supports two assignment strategies: a sliding-window search when the hotspots can be hosted by a single CDR loop, and Hungarian per-residue assignment (Crouse, 2016) when they are distributed across the paratope. Both interventions operate entirely at inference time and require no retraining, making MIMOSA straightforward to integrate with new de novo design models as the field advances. Because it operates on residue-level geometric constraints, MIMOSA is agnostic to antibody format and can, in principle, transfer compatible motifs from any protein–protein interaction.

**Fig. 1:**
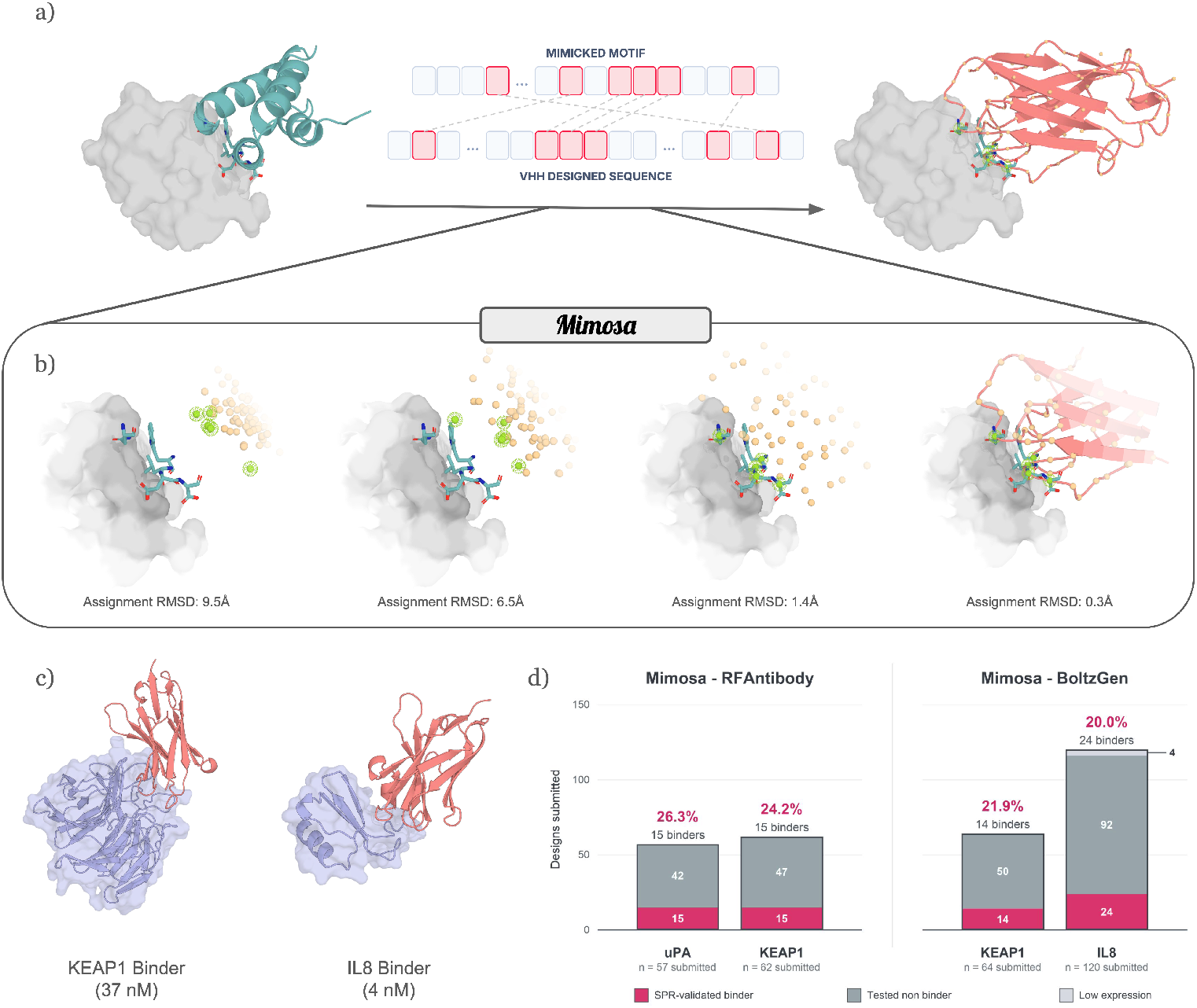
Overview and experimental validation of the MIMOSA framework. **(a)** An example of a mimic antibody, illustrating how the de novo design incorporates critical interaction hotspots from a natural cognate ligand. **(b)** A visual representation of the biasing process during generation, where spatial constraints are imposed at each denoising step on assigned residues, highlighted in green, to guide the model toward binding-competent structural solutions which incorporate the given motif. The Assignment Root Mean Squared Deviation (RMSD) quantifies the spatial deviation between the assigned generated CDR residues and the target motif, and is directly related to the mimic potential (*U*_*m*_) used for steering during generation. **(c)** Computationally designed binders for two targets, KEAP1 and IL-8, with their reported experimental binding affinities of 37 nM and 4 nM, respectively. **(d)** Experimental success ratios of de novo design with mimicry, demonstrating the hit rates achieved when the MIMOSA framework is applied to the RFAntibody and BoltzGen generative pipelines

Here, we validate the framework on single-domain antibodies (VHHs) using both RFAntibody and BoltzGen as generative backbones. We apply it to three targets: uPA and KEAP1, where the motif is placed within a single CDR loop, and interleukin-8 (IL-8), where it is distributed across multiple CDRs. BoltzGen alone was evaluated on two of the three targets and failed to generate binders for either. Applying MIMOSA yielded binders with both generators, with hit rates reaching up to 46% and best affinities of 4 nM against IL-8 and 37 nM against KEAP1 (Fig. 1c,d), unlocking targets that would have been out of reach for the tested de novo methods.

## 2 Results

We validated MIMOSA across three targets and two architecturally independent generators, BoltzGen and RFAntibody (hereafter MIMOSA_BG_ and MIMOSA_RFA_). KEAP1 and uPA are engaged by single contiguous ligand motifs suited to sliding-window mimicry, whereas IL-8 requires per-residue assignment of mimicked residues across multiple CDRs of the designed VHH. This two-dimensional design isolates mimic-guided steering from model-specific behaviour and tests generalisation across interface types.

### 2.1 Mimic-constrained generation yields binders across three targets and two generative models

When unconstrained de novo generation fails on a target, a target-specific prior can help recover it. MIMOSA incorporates this prior at two points in the design pipeline: the mimic potential steers diffusion toward the cognate hotspot geometry, and sequence fixing preserves the corresponding chemistry during inverse folding, as further detailed in the Methods (§4).

Without MIMOSA, BoltzGen did not produce any experimentally validated binder for KEAP1 (0/32) or IL-8 (0/60). By applying our framework, the same model produced binders at hit rates of 22% on KEAP1 (14/64) and 20% on IL-8 (24/120) (Table 1). Applied to a second generator, RFAntibody, the same framework yielded a 24% hit rate on KEAP1 (15/62) and 26% on uPA (15/57). Unconstrained RFAntibody has been reported at a 0–2% hit rate at comparable scale, with individual targets frequently producing zero binders (Bennett et al., 2024), an expected floor that we therefore did not remeasure. All failing targets were recovered, and the recovery was not model-specific.

**Table 1:** Experimental outcomes for the final submitted cohorts. The Template column identifies the interaction template used for mimicry. Percentages in the binder column use all submitted designs as the denominator, including constructs that could not be loaded onto the assay.

| Target | Workflow | Condition | Template | Tested | Binders (%) |
| --- | --- | --- | --- | --- | --- |
| uPA | MIMOSA<sub>RFA</sub> | Geometry-only | 3PB1 | 18 | 7 (38.9%) |
| uPA | MIMOSA<sub>RFA</sub> | Sequence-fixed | 3PB1 | 39 | 8 (20.5%) |
| KEAP1 | MIMOSA<sub>RFA</sub> | Geometry-only | 2FLU | 38 | 4 (10.5%) |
| KEAP1 | MIMOSA<sub>RFA</sub> | Sequence-fixed | 2FLU | 24 | 11 (45.8%) |
| KEAP1 | MIMOSA<sub>BG</sub> | Geometry-only | 2FLU | 32 | 2 (6.3%) |
| KEAP1 | MIMOSA<sub>BG</sub> | Sequence-fixed | 2FLU | 32 | 12 (37.5%) |
| IL-8 | MIMOSA<sub>BG</sub> | Sequence-fixed | 6WZM | 60 | 12 (20.0%) |
| IL-8 | MIMOSA<sub>BG</sub> | Sequence-fixed | 8IC0 | 60 | 12 (20.0%) |
| KEAP1 | BoltzGen | Unconstrained | N/A | 32 | 0 (0.0%) |
| IL-8 | BoltzGen | Unconstrained | N/A | 60 | 0 (0.0%) |

The strongest MIMOSA_BG_ binders reached single-digit nanomolar affinities: a best of 4.1 nM against IL-8 and 37 nM against KEAP1 (Fig. 1c). On IL-8, 23 of 24 fits fell below 100 nM (median 51 nM); the KEAP1 MIMOSA_BG_ binders spanned a wider range (37 nM to 5.1 *µ*M, median 771 nM). MIMOSA_RFA_ binders on KEAP1 and uPA were reported as binary on-target calls without kinetic fits.

### 2.2 Single-loop mimicry: reproducing contiguous ligand motifs in one CDR loop

The simplest form of mimicry is one in which the interaction is driven entirely by a single loop that can be grafted onto a CDR. We tested this mode on two targets where single-loop binders had already been demonstrated (Liu et al., 2017; Kromann-Hansen et al., 2016): KEAP1, engaged by the Nrf2 ETGE motif at the Kelch *β*-propeller (Fig. 2c), and uPA, engaged by the PAI-1 SARMAP loop at the protease cleft (Fig. 2b). Both complexes reported single digit to low double digit nM affinity (Tong et al., 2006; Lo et al., 2006; Eggler et al., 2005; Gong et al., 2016). The mimic potential was applied under the sliding-window assignment strategy (Fig. 2a): the motif was transferred as a single contiguous CDR segment, with the choice of CDR and sequence position left to the generator. Both backbones produced experimentally validated binders at consistent double-digit hit rates: 22% for MIMOSA_BG_ KEAP1 (14/64), 24% for MIMOSA_RFA_ KEAP1 (15/62), and 26% for MIMOSA_RFA_ uPA (15/57) (Table 1). Beyond validating that single-loop mimicry produces binders, this simpler setting also let us evaluate a downstream design choice at inverse folding: whether to fix the recovered hotspot residues to their cognate identity (sequence-fixed) or leave them free for the sequence-design step to choose (geometry-only).

**Fig. 2:**
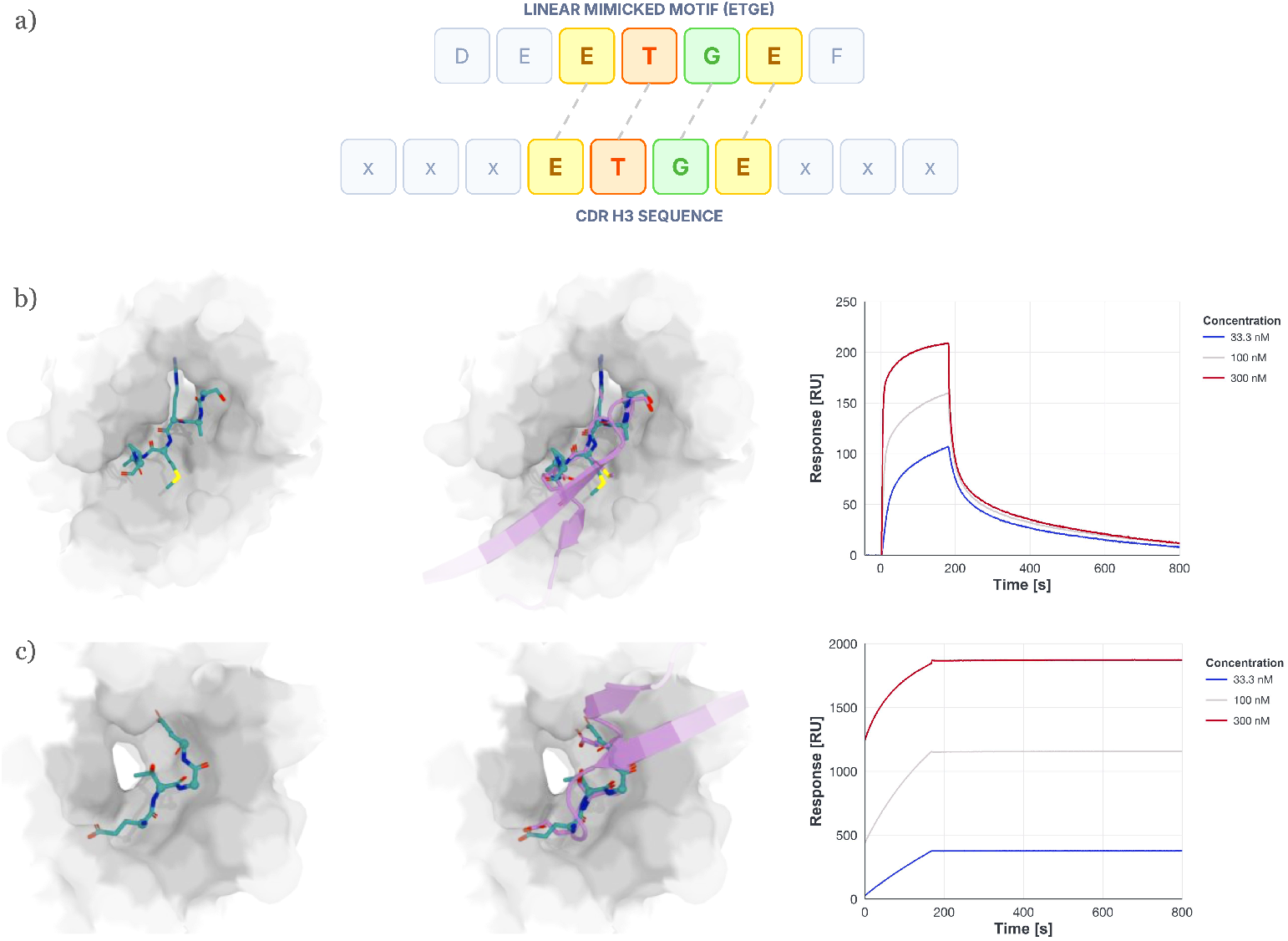
Contiguous single-loop mimicry and experimental validation. **(a)** Schematic of contiguous-motif assignment, illustrated using the Nrf2 ETGE motif. A sliding-window search identifies the CDR segment that best reproduces the ordered motif geometry; matched positions can subsequently be fixed to the cognate residue identities during inverse folding. **(b**,**c)** Representative designs targeting uPA **(b)** and KEAP1 **(c)**. Each row shows the native interaction motif on the target surface (left), its structural reproduction by a CDR of mimic-guided VHH (center), and an SPR sensorgram for the resulting design across increasing target concentrations (right). The uPA design reproduces the PAI-1 SARMAP loop (PDB 3PB1), whereas the KEAP1 design reproduces the Nrf2 ETGE motif (PDB 2FLU).

On uPA, sequence fixing of the whole SARMAP motif did not help binder recovery, and indeed may have slightly hindered it (geometry-only 7/18 vs sequence-fixed 8/39). According to Kromann-Hansen et al. (2016), only one residue in the PAI-1 SARMAP motif is essential for binding to uPA, namely Arg-346, which inserts into the pocket at the base of Asp-189. Their alanine scan of Nb4, a natural nanobody mimic of this interaction, showed the neighbouring positions to be permissive or scaffold-dependent. The physical constraint of the pocket was enough on its own to guide inverse folding, which consistently placed a positively charged residue at the essential position (all 18 geometry-only designs placed Arg or Lys there, as did all seven geometry-only binders, six Arg and one Lys) and successfully designed the surrounding CDR loop to host it. On this target, fixing the whole SARMAP motif therefore added no leverage beyond what inverse folding already provided, and constraining the five permissive positions to PAI-1 identities may have over-constrained designs to a suboptimal solution.

On the other hand, for KEAP1, sequence fixing was essential: hit rate rose from 2/32 (6.3%) to 12/32 (37.5%) for MIMOSA_BG_ and from 4/38 (10.5%) to 11/24 (45.8%) for MIMOSA_RFA_. When we take a closer look at the binders for which the sequence was not fixed, we can identify two glutamates that appear central to binding: the P1 and P4 residues of the Nrf2 ETGE motif were both present in three of these six binders, and at least one was present in all six. Non-binders preserved these residues much less often (6 of 64 with both, 37 of 64 with at least one). This is consistent with what has been observed in the literature: the two glutamates form the salt-bridge core of the interaction with a cluster of Kelch-domain arginines (Tong et al., 2006).

We next explored how sequence fixing affects folding confidence. Refolded with an independent structure model (Protenix-v2 (Zhang et al., 2026)), KEAP1 designs showed a substantial ipSAE (Dunbrack Jr, 2025) lift: median ipSAE was 0.60 for BoltzGen, 0.73 for geometry-only MIMOSA_BG_, and 0.90 for sequence-fixed MIMOSA_BG_; for MIMOSA_RFA_, it rose from 0.46 with geometry-only design to 0.86 with sequence fixing. On uPA, sequence fixing lowered ipSAE for MIMOSA_RFA_ (0.84 geometry-only, 0.70 sequence-fixed). A plausible explanation is that fixing the cognate ETGE motif pushes KEAP1 designs into a geometry the folder already recognises as plausible: the ETGE–Kelch interaction is well represented in the structural training data of natural protein-protein complexes. On uPA, fixing adds no such signal because inverse folding already delivers the essential arginine.

Taken together, the two targets show that sequence fixing helps when an interaction requires multiple cognate residues that inverse folding may struggle to recover on its own, and can be detrimental when the pocket already constrains the essential chemistry or when the motif includes residues that are not critical to binding. With this in mind, we adopt sequence fixing as the default for the scattered-mimicry designs of §2.3.

### 2.3 Scattered mimicry: distributing hotspots across multiple CDRs

The scattered case, in which no single CDR loop can host the whole motif, is the harder mimicry problem. IL-8 illustrates this. It is bound by neutralising antibodies raised against it (e.g. LY3041658, PDB 6WZM) and by its cognate G-protein-coupled receptors (e.g. CXCR1, PDB 8IC0). Both of these binders target the same epitope on IL-8, and do so with overlapping interaction fingerprints.

We applied the framework under the Hungarian per-residue assignment strategy, which allows the mimicked residues to distribute across multiple CDRs of the designed VHH, with no contiguity constraints (Fig. 3a). We used both binder classes as independent templates of interactions to mimic. Each cohort of 60 sequence-fixed MIMOSA_BG_ designs yielded 12 binders (12/58 assayable, 20.7%; Table 1); the two templates were indistinguishable in hit rate, and each independently outperformed the paired unconstrained BoltzGen baseline (0/60). For each template, the mimic motif is a subset of the natural-binder paratope residues whose positions were transferred as the geometric constraint.

**Fig. 3:**
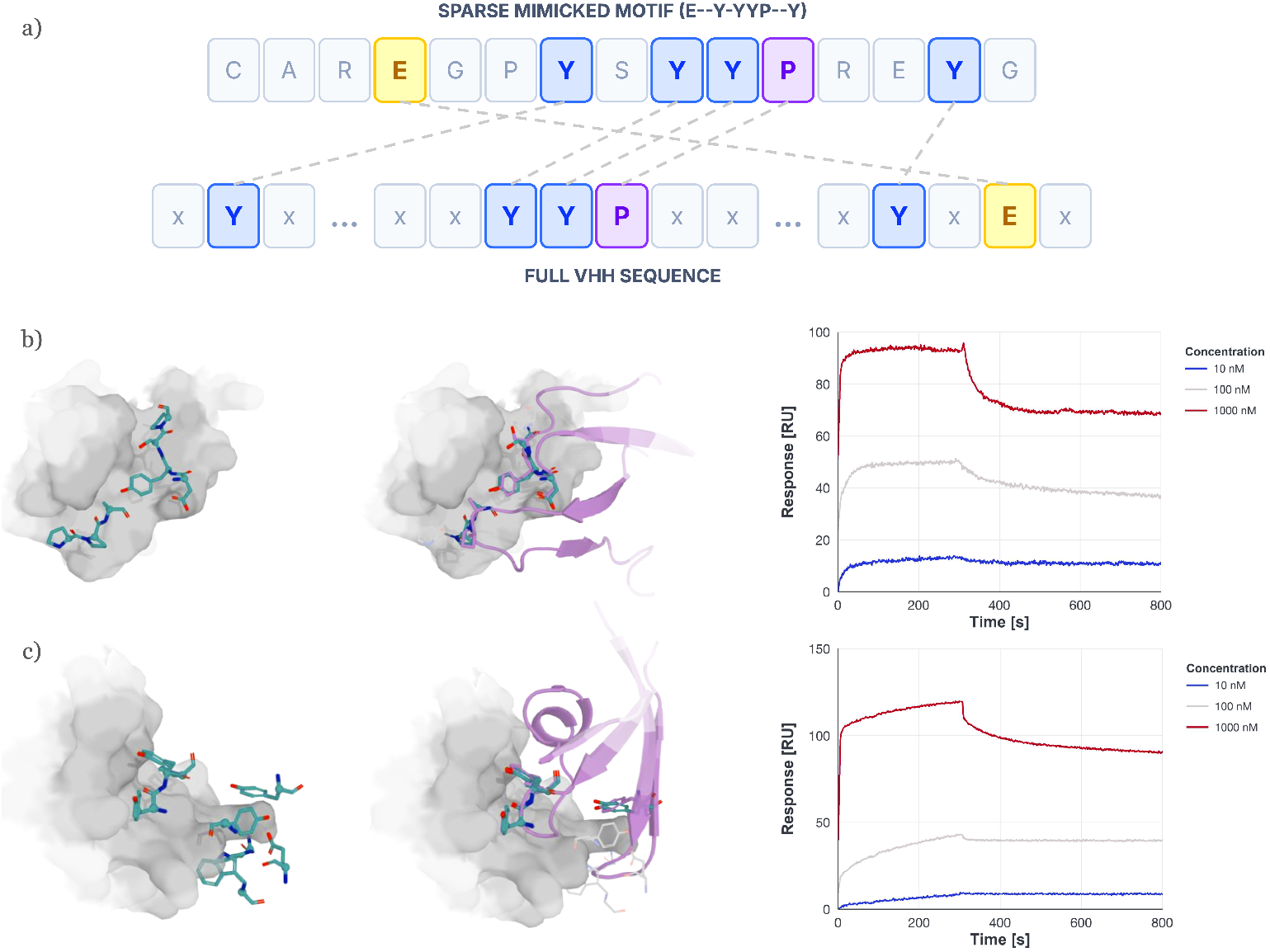
Scattered mimicry across multiple CDR loops and experimental validation. **(a)** Schematic of scattered-hotspot assignment. Non-contiguous residues from the source interface are assigned independently to geometrically compatible positions across the VHH paratope, allowing the interaction pattern to span multiple CDR loops. Matched positions can subsequently be fixed to the corresponding source residue identities during inverse folding. **(b**,**c)** Representative IL-8 designs generated from two independent interaction templates: the receptor-derived CXCR1–IL-8 complex (PDB 8IC0) **(b)** and the antibody-derived LY3041658–IL-8 complex (PDB 6WZM) **(c)**. Each row shows the selected source-interface hotspots on IL-8 (left), their structural reproduction by multiple CDRs of a mimic-guided VHH (center), and an SPR sensorgram for the resulting design across increasing IL-8 concentrations (right).

From LY3041658, we selected seven heavy-chain residues (Trp33, Glu99, Tyr102, Tyr104, Tyr105, Pro106, Tyr110) and one light-chain residue (Trp94) that an alanine scan identified as the key paratope contacts for delivering the Fab’s sub-nM affinity (Boyles et al., 2020). From CXCR1, we selected seven N-terminal residues (Pro21, Pro22, Ala23, Asp26, Tyr27, Ser28, Pro29) that the cryo-EM structure identified as critical for binding (Ishimoto et al., 2023), with the complex having a reported apparent affinity of 4.7nM (Nasser et al., 2009). The selected motifs to be mimicked, their position relative to IL-8, and the CDRs of a representative VHH binder mimicking the motif are shown in Fig. 3b,c. As can be seen, refolding the representative binders from their designed sequences with an independent folding model produces structures whose CDRs reproduce the contacts made by the successfully mimicked paratope residues in the corresponding motif. This suggests that the mimicked interaction geometry is preserved through folding, and is not just an artefact of the steering step.

Across the 24 binders, most of the successfully mimicked residues landed in CDR-H3. Every binder recruited H3, and roughly a third additionally used H2 or H1 as auxiliary loops (Fig. 3b,c). This is consistent with CDR-H3 being longer and more conformationally variable than H1 or H2, allowing it to host arbitrary motif geometries. None of the recovered binders reproduced the entire selected motif, with even the best designs positioning only 3–5 of the 7–8 transferred residues within 2 Å of their target positions. That binders emerged despite this partial mimicry suggests that mimic-constrained generation succeeds by anchoring rather than by fully transplanting the paratope. The mimic potential secures a subset of the interaction geometry, and the generative model completes the interface with de novo contacts that stabilise the complex around the mimicked core.

Folding confidence did not distinguish binder-yielding from non-binder-yielding cohorts on IL-8. Under the same independent Protenix-v2 refolder used for KEAP1, median ipSAE (Dunbrack Jr, 2025) across the three submitted cohorts was similar (0.87 for unconstrained, 0.88 for 6WZM, 0.85 for 8IC0), yet only the mimic-guided cohorts produced binders (0/60 for unconstrained versus 12/60 for each mimic template). At the point of selection, folding-confidence rankings alone would therefore not have differentiated the unconstrained cohort, which failed, from the mimic-guided cohorts, which succeeded, suggesting that mimic quality provides a signal complementary to folding confidence. More analysis on this can be found in the SI.

### 2.4 Structural and sequence determinants of mimic-guided designs

To assess how MIMOSA affects design exploration, we calculated mean normalized pairwise Levenshtein edit distances across IMGT-defined CDR loops. MIMOSA focused CDR sequence diversity, reducing it by up to 20.2% relative to the corresponding target-specific unconstrained BoltzGen cohort. This reduction reflects the intended mechanism: motif-compatible backbone steering during structure generation and hotspot-identity sequence fixing during inverse folding both bias sequences towards motif-hosting conformations. The KEAP1 MIMOSA_BG_ cohort (uniquely carrying all three conditions) decomposed this into an 8.0% reduction from geometry-only MIMOSA_BG_ and a 20.2% reduction from sequence-fixed MIMOSA_BG_ relative to unconstrained BoltzGen. MIMOSA_RFA_ reproduced the effect, with combined-CDR diversity 12.8% lower for KEAP1 and 25.3% lower for uPA relative to geometry-only designs. For contiguous single-loop motifs, sequence fixing produced a strongly CDR-H3-localized contraction (H3 diversity down 23.4% in KEAP1 MIMOSA_BG_, 24.7% in KEAP1 MIMOSA_RFA_, and 45.4% in uPA MIMOSA_RFA_ relative to geometry-only), consistent with the constraint concentrating within the hosting loop. For scattered IL-8 guidance, MIMOSA reduced diversity across multiple CDRs relative to unconstrained BoltzGen (6WZM-guided: 12.0%, 11.6%, and 12.8% across H1, H2, and H3; 8IC0-guided: 19.9% and 13.7% across H2 and H3), consistent with scattered assignment allowing multiple loops to contribute. Despite this focusing of sequence space, every submitted combined-CDR sequence remained unique, and MIMOSA_BG_ designs still differed by an average of 53%–61% across their CDRs, compared with 66%–69% in unconstrained cohorts.

Beyond focusing sequence space, MIMOSA also shifts which framework scaffolds and CDR loop lengths are selected. From a shared set of five VHH frameworks for IL-8 (7eow, 8coh, 3eak, 8z8v, 7xl0), unconstrained BoltzGen concentrated on a single scaffold (7eow, 47 of 60 submissions), whilst MIMOSA_BG_ redistributed toward 8coh (56 of 120) and 3eak (41 of 120), frameworks on which guidance more often succeeded in placing the motif. MIMOSA did not similarly narrow loop-length exploration. The 120 IL-8 MIMOSA submissions occupied 82 distinct IMGT CDR-length combinations, with the 24 recovered binders spanning 18 combinations and three frameworks; sequence-fixed KEAP1 MIMOSA_BG_ designs likewise sampled 32 distinct combinations across 32 submissions. For MIMOSA_RFA_, whose sampler covers fewer H3 lengths overall, KEAP1 binder recovery was strongly concentrated at IMGT CDR-H3 length 12 (12 of 15 confirmed binders, 12 of 20 tested at that length versus 3 of 42 at all other lengths), whereas uPA binders were recovered at two of the three submitted lengths. Together, these results suggest that MIMOSA recovers binders across multiple scaffold–loop configurations, adapting to the preferences of each target and generator.

At the residue level, the de novo design model plays two complementary roles, introducing additional contacts beyond the mimicked motif and shaping the CDR to present it at the productive geometry. Both are visible in Protenix-v2 refolds of the KEAP1 MIMOSA_RFA_ cohort. On the Kelch surface, 14 of 15 binders (versus 21 of 47 non-binders) engaged Asp205 through the C-terminal residue of CDR-H2, in ten cases as a lysine or arginine forming a salt bridge. Asp205 sits at the periphery of the ETGE binding pocket and is not engaged by Nrf2 in the native complex (PDB 2FLU); it is an additional contact site the design model recruited alongside the mimicked motif. Within CDR-H3 itself, 9 of 11 binders that preserved the ETGE motif (versus 2 of 13 non-binders) carried a glycine three positions upstream of the motif. The glycine never contacts KEAP1 in any binder structure, consistent with a role in shaping the CDR-H3 loop that presents the ETGE motif at the productive geometry.

## 3 Discussion

MIMOSA is a model-agnostic, inference-time framework that helps recover targets on which unconstrained de novo antibody design produces zero or near-zero binders. Applied without retraining to two architecturally distinct diffusion-based generators (RFAntibody and BoltzGen (Bennett et al., 2024; Stark et al., 2025)), it delivered hit rates of 20–26% on all three targets tested: KEAP1, IL-8, and uPA. The unconstrained BoltzGen baseline produced no binders for KEAP1 and IL-8, and unconstrained RFAntibody has been reported at 0–2% (Bennett et al., 2024). The two mimicking strategies introduced here, sliding-window and Hungarian per-residue assignment, let the framework place the mimic motif either as a contiguous CDR insert or across multiple CDRs of the designed VHH.

The design-prior interpretation of these results is straightforward. Cognate ligand motifs represent experimentally validated solutions to the binding problem, defined at the level of residue identity, backbone geometry and interaction chemistry. Constraining a generative model to reproduce these features injects a proven interaction blueprint into the search, biasing it toward regions of design space known to yield productive binders. Immune repertoires appear to converge on the same principle: naturally occurring antibodies against these and related targets mimic cognate ligand motifs, either within a single CDR loop or across the paratope (Kromann-Hansen et al., 2016; Abanades et al., 2026). Prior work exploited this convergence computationally by grafting known cognate motifs onto natural antibody scaffolds (Liu et al., 2017); the framework introduced here generalises that idea by constraining a generative model to construct scaffolds compatible with the cognate binding motif at inference time, and by extending the approach to scattered assignment patterns. MIMOSA exploits an asymmetry that unconstrained methods do not: when a cognate binder is known, part of the binding problem has already been solved.

This constraint focuses designed sequence pools without collapsing them onto a small number of solutions. Combined-CDR diversity decreased by up to 20.2% relative to unconstrained cohorts, with sequence fixing producing the strongest contraction in the motif-hosting loops. Nevertheless, every submitted combined-CDR sequence remained unique, and guided designs still differed from one another by an average of 53–61% across their CDRs. This focusing was also compatible with broad structural exploration: binders were recovered across multiple framework and loop-length configurations. MIMOSA therefore acts as a targeted restriction of the search space to binding-competent designs rather than as a prescription of a single antibody solution.

One way MIMOSA can complement current de novo methods is at the inverse-folding step, though whether it helps or hinders depends on which residues are selected for the mimic motif. On KEAP1, where the ETGE motif contains two essential glutamates, fixing the motif dramatically improved binder recovery, raising MIMOSA_BG_ hit rate from 2/32 to 12/32 and MIMOSA_RFA_ hit rate from 4/38 to 11/24, with every geometry-only MIMOSA binder recovering at least one of the two critical glutamates. On uPA, where the six-residue SARMAP motif contains only one residue essential for binding (Arg-346), fixing the whole motif did not help and even slightly hindered recovery (8/39 sequence-fixed vs 7/18 geometry-only). All seven geometry-only binders independently recovered the essential positively charged residue (six Arg and one Lys), showing that the key interaction could be preserved without fixing the whole motif. Choosing which motif residues to constrain, preserving the essential interaction while retaining the generator’s freedom to optimise the surrounding sequence, is therefore a critical design decision, likely highly dependent on the target and cognate ligand being mimicked. The optimal constraint will often not be the whole cognate motif but the minimal fingerprint sufficient to anchor productive recognition, leaving the generator free to construct the surrounding paratope.

Another way MIMOSA can complement current de novo methods is at the design-selection step. Folding confidence and mimic fidelity are complementary signals available for ranking or filtering designs: one measures how confident an independent structure model is in the predicted interface, the other how faithfully a design reproduces a known interaction fingerprint. On KEAP1, the two were strongly correlated and tracked binder recovery similarly, with median ipSAE increasing progressively from 0.60 for BoltzGen to 0.73 for geometry-only MIMOSA_BG_ and 0.90 for sequence-fixed MIMOSA_BG_. On IL-8 they decoupled. All three submitted cohorts showed similar folding confidence (ipSAE 0.85–0.88), yet only the mimic-guided cohorts produced binders (0/60 for unconstrained versus 12/60 for each mimic template). Folding confidence would not have discriminated between these cohorts, whereas mimic fidelity did. Filtering mimic-guided designs on both metrics is therefore likely more robust than folding confidence alone. Folding confidence flags globally implausible structures, and mimic fidelity preserves binding-relevant geometry. This complementarity matters most on targets where the two decouple.

The three targets studied here also begin to define the interface properties for which MIMOSA is likely to be most effective. KEAP1, uPA, and IL-8 all contain well-characterised, high-affinity interaction fingerprints in which a limited number of residues provide strong geometric and chemical constraints. KEAP1 represents an interaction dominated by a small number of chemically essential residues, uPA an even more minimal anchor centred on a single critical residue, and IL-8 a fingerprint distributed across multiple CDR loops of the designed antibodies. Together, these examples suggest MIMOSA is particularly well suited to targets whose binding geometry is defined by a compact set of interaction hotspots. Notably, MIMOSA does not require exact transplantation of the natural interface. On uPA, a single essential residue anchored the designed binders, and on IL-8, binders emerged despite reproducing only subsets of the transferred fingerprint. In both cases, a relatively sparse geometric anchor sufficed for the generative model to complete a productive interface.

These strengths also define the current boundaries of the approach. MIMOSA presently requires a structurally characterized target–binder complex from which a transferable interaction fingerprint can be extracted; whether sufficiently accurate predicted complexes can substitute for experimental structures remains to be established. Interfaces that are large or energetically diffuse, with many residues making individually modest contributions, may provide less effective guidance because no compact fingerprint can be readily identified. Because the source structure is static, strongly conformational or induced-fit interfaces may not be adequately captured by a single template. Finally, MIMOSA inherits the epitope of the source interaction, which is advantageous when reproducing a known functional binding site but restrictive when an arbitrary or novel epitope is desired.

These limitations define a complementary rather than universal role for mimic-guided design. Unconstrained generation remains essential when no interaction prior exists or when novel epitope discovery is the objective. When a functional interaction is already structurally characterised, however, ignoring that information forces a generative model to rediscover a binding solution that is already known. De novo antibody design ultimately faces two problems: identifying an interaction geometry capable of supporting binding and constructing an antibody that realises it. Current generative approaches attempt to solve both simultaneously. MIMOSA separates them by borrowing a validated interaction fingerprint from the cognate binder and leaving the generative model to construct the surrounding paratope. As de novo antibody models become increasingly capable, such interaction priors offer a complementary route toward more reliable binder discovery across challenging targets.

## 4 Methods

Diffusion-based *de novo* antibody-design workflows typically comprise three stages: generation of a candidate antibody-target structure, inverse folding to assign or refine the antibody sequence, and refolding-based validation of the designed complex. Both BoltzGen and RFAntibody follow this general workflow, but use different molecular representations during diffusion (Bennett et al., 2024; Stark et al., 2025). BoltzGen generates complexes using an all-atom representation, whereas RFAntibody generates antibody backbones using residue-level rigid frames. This distinction makes the two systems useful tests of whether the same geometric guidance strategy can transfer across generator architectures

MIMOSA interfaces at two stages of this workflow. During structure generation, a differentiable mimic potential (*U*_*m*_) steers diffusion toward backbones that reproduce the geometry of a reference interaction motif. During inverse folding, the identities of successfully placed motif residues can be fixed while the remaining CDR sequence is designed. The following sections describe these two interventions in turn.

### 4.1 Mimic-constrained generation

We implemented inference-time steering within two diffusion-based antibody generation frameworks, RFAntibody and BoltzGen (Bennett et al., 2024; Stark et al., 2025), to guide sampling trajectories towards designs that reproduce the interaction we want to mimic. At each denoising step, the predicted structure is corrected by gradient descent on a differentiable mimic potential *U*_*m*_, following the backward universal guidance framework (Bansal et al., 2023). To avoid entrapment in local minima, we additionally employed Feynman–Kac particle steering (Singhal et al., 2025; Hartman et al., 2025), evolving multiple parallel trajectories and resampling in favour of low-energy states. This setup is largely inspired by the one presented in Boltz-Steering (Wohlwend et al., 2025). This guidance procedure was applied to both generative models, as it is model architecture-agnostic and can be similarly applied to any diffusion-based framework.

#### Mimic Potential

We introduce the mimic potential *U*_*m*_, which quantifies how well the generated CDR reproduces the spatial arrangement of the mimicked motif. The calculation fundamentally consists of three distinct steps: alignment, assignment and computation. The alignment phase ensures the motif coordinates are in the same frame of reference as the predicted target, by applying the same transformation that rigidly superimposes the reference target onto the predicted target. During assignment we identify the optimal correspondence *π*^*∗*^ between CDR residues and motif positions which minimizes the sum of distances. When mimicking contiguous motifs, a sliding-window search identifies the best-matching CDR segment; for general scattered motifs, the Hungarian algorithm solves the assignment exactly (Crouse, 2016). The potential is then computed as:

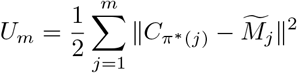

where *m* is the number of motif residues, 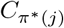 are the assigned C*α* coordinates, and 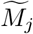 the aligned motif positions. More details on the implementation are provided in the SI.

### 4.2 Sequence fixing during inverse folding

The mimic potential spatially guides structure generation but does not enforce residue identity. Although RFAntibody generates backbone coordinates while BoltzGen generates full-atom structure with a complete sequence, both workflows follow a framework comprising structure generation, inverse folding, and refolding. To preserve the interaction chemistry of the mimicked motif, we intervene in the inverse folding step by fixing the sequence of successfully placed mimic residues. Specifically, after generation, alignment and assignment are re-evaluated on the final structure; CDR residues whose C*α* atoms fall within 2 Å of their assigned motif position are fixed to the respective amino acid identity. The remaining CDR positions are then repredicted by the inverse folding model (ProteinMPNN (Dauparas et al., 2022) for RFAntibody; BoltzIF for BoltzGen), conditioned on the fixed mimic residues, framework, and antigen.

### 4.3 Expression

VHH designs were expressed using three strategies matched to the downstream SPR capture chemistries. Mammalian expression was performed in-house at Roche: constructs bearing a C-terminal Avitag (for *in vivo* biotinylation) and C-tag (for purification) were transiently transfected into Expi293 cells (Thermo Fisher) and purified by automated CaptureSelect C-tag affinity chromatography, with an optional size-exclusion polish. Cell-free expression was performed at Adaptyv Bio: constructs carrying a C-terminal twin-strep affinity tag were produced from pre-assembled genes in a prokaryotic in-vitro translation system, with concentration determined via an affinity-based quantification assay. *E. coli* expression was performed in-house at Roche: constructs bearing C-terminal V5 and His tags were purified by His-tag affinity chromatography. All purified products were quality-controlled by capillary electrophoresis SDS (CE-SDS).

### 4.4 Binding characterisation

Binding was measured by surface plasmon resonance (SPR) on Carterra platforms. VHHs were captured via their C-terminal affinity tag (biotinylated ligands on streptavidin-functionalised chips; twin-strep-tag ligands on Strep-Tactin XT-functionalised chips; V5-tag ligands on sensor chips functionalised with a mouse IgG1 anti-V5 antibody (Chromotek)), and recombinant human KEAP1, uPA, and IL-8 were injected as analytes in a single-cycle kinetics format over a graded concentration series in HBS-EP+ running buffer. Sensorgrams were double-referenced and globally fitted to extract apparent kinetic parameters. Designs were classified as binders when a *K*_*D*_ could be reliably fitted, or when the association-phase signal exceeded the negative control by at least 300% without a fit. Constructs that could not be loaded onto the sensor were counted with the non-binders for the reported hit rates. Reported *K*_*D*_ values are apparent, as fits were predominantly bivalent-analyte rather than 1:1. More details on expression and binding are given in the SI.

