## Supplementary Information for "MIMOSA: Guiding De Novo Antibody Design with Natural Interaction Fingerprints"

### A. Supplementary Methods

#### A1. Detailed formulation of the mimic potential

The mimic potential $U_m$ is the mechanism through which MIMOSA constrains antibody diffusion to reproduce a chosen cognate interaction. It is a differentiable coordinate-space score that quantifies the geometric agreement between generated CDR residues and a set of source-motif residues drawn from a cognate ligand–target complex. The correspondence between design and source-motif residues is not fixed a priori; it is resolved by a distance-minimizing assignment each time the potential is evaluated.

##### A1.1 Target-frame alignment

The alignment maps the source-motif coordinates from their native frame into the frame of the current denoised prediction. Their native frame is the cognate ligand–target complex, from which the target and the source-motif residues are retained. Let $M_j \in \mathbb{R}^3$ denote the $C\alpha$ coordinate of source-motif residue $j$; $R_k \in \mathbb{R}^3$, $k = 1, \dots, K$, the $C\alpha$ coordinates of the reference target; and $X_k \in \mathbb{R}^3$ the $C\alpha$ coordinates of the corresponding target residues in the prediction. Correspondence between $R_k$ and $X_k$ is established by pairwise sequence alignment when the two target sequences differ. At each evaluation, the rigid transformation aligning the reference target to the prediction is obtained by the Kabsch solution

$$(Q^*, b^*) = \arg \min_{Q \in SO(3), b \in \mathbb{R}^3} \sum_{k=1}^K \|QR_k + b - X_k\|_2^2 \quad (\text{S1})$$

and applied to each source-motif coordinate:

$$\widetilde{M}_j = Q^* M_j + b^*. \quad (\text{S2})$$

Recomputing the alignment at each evaluation makes the potential invariant to global translation and rotation of the generated complex and constrains the motif relative to the target.

##### A1.2 Dynamic residue assignment

Let $C_i$, $i = 1, \dots, N$, be the ordered $C\alpha$ coordinates of the eligible design residues (the CDR positions that may host the motif) and let $m$ be the number of source-motif residues. The pairwise assignment cost is the Euclidean distance

$$d_{ij} = \|C_i - \widetilde{M}_j\|_2. \quad (\text{S3})$$

The assignment minimizes the sum of these distances, whereas the energy applied after assignment is the half-sum of their squares.

For the ordered KEAP1 and uPA motifs, the contiguous assignment preserves motif order and searches over sliding windows:

$$a^* = \arg \min_{a \in \{1, \dots, N-m+1\}} \sum_{j=1}^m d_{a+j-1, j}, \quad \pi^*(j) = a^* + j - 1. \quad (\text{S4})$$

For the scattered IL-8 interfaces, correspondence is an injective linear assignment with no sequence-adjacency constraint:

$$\pi^* = \arg \min_{\pi \text{ injective}} \sum_{j=1}^m d_{\pi(j), j}, \quad (\text{S5})$$

solved by the rectangular Hungarian algorithm [1]. This lets different source-motif residues be placed across different CDRs.

The target alignment and selected assignment are treated as stop-gradient operations during an individual update, and both are recomputed whenever the mimic potential is evaluated.

#### A1.3 Energy and gradient

Given the selected correspondence, the mimic potential that biases the design toward the cognate interaction is defined as

$$U_m(C; M) = \frac{1}{2} \sum_{j=1}^m \left\| C_{\pi^*(j)} - \widetilde{M}_j \right\|_2^2. \quad (\text{S6})$$

With the alignment and assignment held fixed during an individual update, its $\text{C}\alpha$ gradient is

$$\nabla_{C_i} U_m = \begin{cases} C_i - \widetilde{M}_j, & i = \pi^*(j), \\ \mathbf{0}, & i \notin \text{im}(\pi^*). \end{cases} \quad (\text{S7})$$

### A2. Steering of the diffusion process

MIMOSA modifies sampling at inference time and requires no retraining of the underlying generator. The mimic potential $U_m$ (Section A1) is minimized during reverse diffusion by combining backward guidance with Feynman–Kac [2] particle resampling, following the strategy of Boltz-steering [3]. We apply the same guidance procedure to two structurally distinct reverse-diffusion processes: the residue-frame representation of RFAntibody and the all-atom representation of BoltzGen [4, 5]. We refer to the corresponding implementations as MIMOSA<sub>RFA</sub> and MIMOSA<sub>BG</sub>, respectively. At reverse step $i$, let $z_i$ be the current noisy state, $\tau_i$ the generator-specific diffusion time or noise parameter, and $\widehat{x}_{0,i} = D_\theta(z_i, \tau_i)$ the model’s current estimate of the clean structure. The mimic potential $U_m(\widehat{x}_{0,i})$ is evaluated on this denoised estimate rather than directly on $z_i$, and the resulting coordinate update is then incorporated into the generator-specific reverse transition.

#### A2.1 Backward guidance

We perform gradient guidance using the backward universal guidance framework [6], applying one gradient update at each active, non-terminal step. Let $g_{i,r} = \nabla_{C_r} U_m$ be the mimic-potential gradient calculated at the $\text{C}\alpha$ coordinate of design residue $r$. Both generators use the same residue-level gradient but apply it through their respective structural representations.

RFAntibody represents residue geometry as frames centred on the $\text{C}\alpha$ atoms. If $(R_{i,r}, t_{i,r})$ denotes the rotation and translation of residue frame $r$, guidance translates the frame origin:

$$\mathcal{G}_{i,r}(R_{i,r}, t_{i,r}) = (R_{i,r}, t_{i,r} - \eta_i g_{i,r}). \quad (\text{S8})$$

Guidance therefore acts only on the translational diffusion process; the IGSO(3) rotational process is unchanged.

BoltzGen represents atoms directly in three-dimensional Cartesian coordinates. The same $\text{C}\alpha$-derived gradient is applied unchanged to every represented atom $a$ of residue $r$:

$$\widetilde{x}_{0,i,r,a} = \widehat{x}_{0,i,r,a} - \eta_i g_{i,r}. \quad (\text{S9})$$

Each update is therefore a rigid translation of the residue. This is geometrically equivalent to RFAntibody’s frame-origin translation, despite the different underlying representations.

We use a cosine schedule for the guidance coefficient $\eta_i$, ramping from 0 at $s_i = 0$ (the noisiest reverse step) to a maximum of 0.3 at $s_i = 1$ (the final denoised step). The ramp starts at zero to reduce sensitivity to uncertainty in early denoised estimates, particularly in the exact residue-level assignment of the motif. As these estimates stabilize, the smooth rise progressively strengthens the guidance without abrupt changes in guidance strength.

$$\eta_i = \frac{0.3}{2} [1 - \cos(\pi s_i)]. \quad (\text{S10})$$

The gentle ramp is not sufficient by itself. Because the update magnitude $\eta_i \|g_{i,r}\|$ scales with the gradient norm, the coefficient ceiling of 0.3 is not a 0.3-Å displacement cap. Large gradients can produce independent translations of adjacent residues that overpower the learned reverse dynamics and disrupt peptide-bond geometry. We therefore combine the ramp with the post-sampling backbone-integrity filter described in Section A5.2.

### A2.2 Feynman–Kac particle steering

Backward guidance locally changes each trajectory but can still lead a single trajectory into an unfavorable basin. We therefore combine it with Feynman–Kac (FK) steering [2], following the particle-resampling construction used by Boltz–Steering [3]. Four particles are advanced in parallel, each initialized from an independent draw of the noise prior. At resampling event $\ell$, let $E_\ell^{(k)} = U_m(\hat{x}_{0,\ell}^{(k)})$ be the resulting energy of particle $k$, evaluated on the unguided denoised estimate. The incremental FK term is

$$\log G_\ell^{(k)} = \begin{cases} -E_0^{(k)}, & \ell = 0, \\ E_{\ell-1}^{(A_{\ell-1}(k))} - E_\ell^{(k)}, & \ell > 0, \end{cases} \quad (\text{S11})$$

where $A_{\ell-1}(k)$ is the ancestor of particle $k$ at the preceding resampling event. The unnormalized log weight is

$$\log w_\ell^{(k)} = \lambda \log G_\ell^{(k)} + \Delta_\ell^{(k)}. \quad (\text{S12})$$

Here $\lambda$ controls selection strength, and $\Delta_\ell^{(k)}$ is the generator-specific proposal correction accumulated since the preceding resampling event. Normalized weights are obtained with a softmax, and particles are resampled multinomially with replacement every three reverse steps. At the terminal event, one particle was sampled stochastically from the normalized weights.

Table S1: Steering settings shared by MIMOSA<sub>RFA</sub> and MIMOSA<sub>BG</sub>. The cosine schedule is written against noisy-to-clean denoising progress, as defined in Eq. S10.

| Parameter | Value |
| --- | --- |
| Gradient updates per active reverse step | 1 |
| Backward-guidance schedule | cosine, $0 \rightarrow 0.3$ |
| FK particles | 4 |
| FK resampling interval | every 3 reverse steps |
| FK potential coefficient, $\lambda$ | 4.0 |

Because guidance changes the proposal, the FK weight must include a likelihood-ratio correction. Let $K_i$ denote a generator’s native unguided reverse-transition kernel and $Q_i$ the guided proposal at reverse step $i$. For the set $\mathcal{I}_\ell$ of guided steps since the preceding resampling event, the correction in Eq. S12 is

$$\Delta_\ell^{(k)} = \sum_{i \in \mathcal{I}_\ell} \log \frac{K_i(z_{i-1}^{(k)} | z_i^{(k)})}{Q_i(z_{i-1}^{(k)} | z_i^{(k)})}. \quad (\text{S13})$$

The native kernels differ between the two generators, but MIMOSA changes only Gaussian translational components in both. If $K_i(y | z) = \mathcal{N}(\mu_i, v_i I)$, $Q_i(y | z) = \mathcal{N}(\mu_i + \delta_i, v_i I)$, and $y = \mu_i + \delta_i + \epsilon_i$ is a guided sample, the per-step correction has the shared mean-shift form

$$\log \frac{K_i(y | z)}{Q_i(y | z)} = \frac{\|\epsilon_i\|_2^2 - \|\epsilon_i + \delta_i\|_2^2}{2v_i}. \quad (\text{S14})$$

For RFAntibody, this ratio is evaluated over one three-dimensional translation per diffused residue frame. Its rotational kernel is unchanged by guidance, so the corresponding IGSO(3) ratio is one and contributes zero to the log correction. For BoltzGen, the ratio is evaluated in the affected all-atom Cartesian coordinate space. The two generators therefore use different kernel-specific displacements and variances, while sharing the Gaussian mean-shift structure.

The likelihood-ratio treatment used here differs from that shown in Algorithm 2 of the Boltz-1 manuscript [3]. In that algorithm, guidance is applied at every reverse step while resampling occurs every three steps, and the resampling weight includes only the likelihood ratio for the transition at the resampling checkpoint. To implement the SMC formulation of FK steering more formally, we instead accumulate the ratios for all guided transitions in the interval between successive resampling events, as in Eq. S13.

#### A3. Interaction templates and biological context

Table S2: Source interactions used to define the mimic constraints.

| Target | Source structure | Source partner | Assignment topology |
| --- | --- | --- | --- |
| KEAP1 | PDB 2FLU | Nrf2 **ETGE** turn | Ordered contiguous window |
| uPA | PDB 3PB1 | PAI-1 **SARMAP** loop | Ordered contiguous window |
| IL-8 | PDB 6WZM | LY3041658 antibody interface | Scattered one-to-one assignment |
| IL-8 | PDB 8IC0 | CXCR1 receptor interface | Scattered one-to-one assignment |

##### A3.1 Contiguous interaction templates

KEAP1 is an adaptor of the Cullin-3 ubiquitin-ligase complex that regulates the oxidative-stress response by recruiting Nrf2 for degradation. Nrf2 binds the Kelch β-propeller through its high-affinity **ETGE** degron, whose two glutamate side chains form the electrostatic core of the interaction with an arginine-rich pocket in KEAP1 [7, 8]. We use the ordered **ETGE** turn from the KEAP1–Nrf2 complex (PDB 2FLU), an interaction for which hotspot grafting has previously produced a designed antibody binder [9].

uPA is a serine protease in the plasminogen-activation system. Its inhibitor PAI-1 presents a reactive-centre loop that occupies the protease cleft in a substrate-like geometry. We use the ordered **SARMAP** segment from the uPA–PAI-1 Michaelis complex (PDB 3PB1). Within this segment, Arg-346 inserts into the uPA specificity pocket and contacts Asp-189 at the base of the pocket. The natural camelid VHH Nb4 reproduces this binding mode with its CDR-H3, and alanine scanning identified Arg-346 as the sole essential position within the **SARMAP** segment [10]. The KEAP1 and uPA templates therefore test transfer of biologically established, ordered single-loop recognition motifs.

##### A3.2 Scattered interaction templates

IL-8 is a chemokine recognized both by neutralizing antibodies and by its cognate chemokine receptors. We derived two independent scattered templates from interactions that partially overlap on the IL-8 surface but originate from different partner classes. The antibody-derived template uses seven LY3041658 heavy-chain residues (Trp33, Glu99, Tyr102, Tyr104, Tyr105, Pro106, and Tyr110) and the light-chain residue Trp94 from the antibody–IL-8 interface (PDB 6WZM), identified as key paratope contacts in the crystal structure of Boyles et al. [11]. The receptor-derived template uses seven residues from the CXCR1 N terminus (Pro21, Pro22, Ala23, Asp26, Tyr27, Ser28, and Pro29) at the receptor–IL-8 interface (PDB 8IC0), identified as critical for binding by the cryo-EM structure of Ishimoto et al. [12]. Unlike the contiguous templates, these interaction residues are assigned independently and can be distributed across multiple VHH CDRs.

#### A4. Sequence fixing

The geometric guidance does not by itself require an assigned design residue to retain the amino-acid identity of its source residue. We therefore compare two inverse-folding strategies. In the *geometry-only* strategy, the generated backbone is passed to inverse folding without fixing the source identities.

In the *sequence-fixed* strategy, target alignment and residue assignment are re-evaluated on the final generated structure. Assigned pair $j \mapsto \pi^*(j)$ is fixed only when

$$\left\| C_{\pi^*(j)} - \widetilde{M}_j \right\|_2 < 2.0 \text{ \AA}. \quad (\text{S15})$$

Each passing design position is set to the amino-acid identity of the corresponding source residue and given as additional context to the inverse folder. Assigned positions outside the threshold remained designable. The remaining designable positions are then predicted by the inverse-folding model (ProteinMPNN [13] for RFAntibody, BoltzIF for BoltzGen), conditioned on the fixed mimic residues, framework, and antigen sequences.

### A5. Sampling, filtering, and selection

The computational campaigns were conducted over several months, during which de novo models, structure-prediction models, and our filtering and selection strategies evolved. The procedures were therefore not identical across campaigns: each used the models and criteria that, at the time, reflected the available state of the art. We underline, however, that for a given target and de novo model (eg, unconstrained BoltzGen, geometry-only MIMOSA<sub>BG</sub>, and sequence-fixed MIMOSA<sub>BG</sub> against KEAP1), strategies were consistent and thus directly comparable.

#### A5.1 Input configurations

For the KEAP1 and uPA MIMOSA<sub>RFA</sub> campaigns, we followed the example configuration in the RFAntibody repository, which uses the **3eak**/h-NbBCII10 VHH framework and a Chothia-based loop convention [4]. We retained its design lengths $H1 = 7$ and $H2 = 6$ and extended only the allowed $H3$ range, from 5–13 to 5–17. In the IMGT numbering scheme used in Section B, lab-submitted MIMOSA<sub>RFA</sub> designs had $H1 = 8$, $H2 = 8$, and $H3$ lengths spanning 10–19.

All BoltzGen and MIMOSA<sub>BG</sub> campaigns used the variable loop length definitions supplied with the original BoltzGen nanobody scaffold configurations for **7eow**, **7xl0**, **8coh**, and **8z8v** [5], together with the RFAntibody **3eak** scaffold and its corresponding framework and CDR design spans. Thus, each of these campaigns sampled from the same five-framework menu: **3eak**, **7eow**, **7xl0**, **8coh**, and **8z8v**. The final lab-submitted framework distributions are outcomes of generation, filtering, and selection.

Beyond the framework choices and other deviations explicitly described here, all input and generator settings were retained at their repository defaults.

#### A5.2 MIMOSA-specific structural checks

In addition to the standard generator filters, MIMOSA candidates were subject to two structural checks that were not imposed on the corresponding unconstrained cohorts. First, because gradient guidance can distort local backbone geometry when its updates become too strong, generated candidates were checked for peptide-bond integrity before downstream scoring. Within each chain, consecutive residues $r$ and $r + 1$ were required to satisfy

$$\left| \|C_r - N_{r+1}\|_2 - 1.329 \text{ \AA} \right| \leq 0.5 \text{ \AA}. \quad (\text{S16})$$

Any candidate containing a consecutive-residue pair outside this tolerance was discarded, avoiding inverse folding and refolding steps.

Second, after target-frame alignment and residue assignment as described in Sections A1.1 and A1.2, mimic motif recovery was quantified by the Cα RMSD over the selected correspondence:

$$\text{RMSD}_{\text{motif}} = \left( \frac{1}{m} \sum_{j=1}^m \left\| C_{\pi^*(j)} - \widetilde{M}_j \right\|_2^2 \right)^{1/2}. \quad (\text{S17})$$

This quantity was evaluated on the relevant generated backbone or refolded structure as specified for each campaign below; the applicable thresholds were campaign-specific.

#### A5.3 MIMOSA<sub>RFA</sub> KEAP1 and uPA sampling, filtering, and selection

For each target, we generated 2,000 backbones with MIMOSA<sub>RFA</sub> guidance and retained those with a motif-recovery RMSD (Eq. S17) below 2.0 Å. To preserve diversity in the orientation of the VHH relative to the target, each passing backbone was represented by the Cα coordinates of the first and last eight VHH residues after target alignment. Pairwise RMSDs over these 16 terminal framework residues provide an approximate measure of binding pose. Backbones were greedily clustered using this measure with a 6.0 Å cutoff, and 32 backbones per target were selected by round-robin sampling across clusters.

For each selected backbone, ProteinMPNN generated 100 sequences under two conditions: geometry-only MIMOSA<sub>RFA</sub> and sequence-fixed MIMOSA<sub>RFA</sub>. Each sequence was refolded with RF2 and independently predicted and scored with Boltz-2 [14]. We required all three pairwise target-aligned CDR RMSDs (Boltz-2 versus RF2, Boltz-2 versus the generated backbone, and RF2 versus the generated backbone) to be below 2.0 Å. Duplicate VHH sequences were removed. The three-way self-consistency selection yielded 216 KEAP1 and 505 uPA designs. Passing candidates were ranked by Boltz-2 iPTM, newly clustered by structural diversity on the Boltz-2 predicted structure, and selected by round-robin sampling across clusters.

The submitted cohorts contained 62 KEAP1 designs (38 geometry-only and 24 sequence-fixed) and 57 uPA designs (18 geometry-only and 39 sequence-fixed).

#### A5.4 BoltzGen and MIMOSA<sub>BG</sub> KEAP1 sampling, filtering, and selection

For KEAP1, 60,000 designs were generated under each of three protocols: unconstrained BoltzGen, geometry-only MIMOSA<sub>BG</sub>, and sequence-fixed MIMOSA<sub>BG</sub>. Following repository defaults, each backbone was inverse folded once and refolded. Candidates were required to pass the standard BoltzGen sequence and structure filters. For the unconstrained cohort, the top 300 designs were retained according to the native BoltzGen composite quality ranking. For each MIMOSA<sub>BG</sub> cohort, the motif-recovery RMSD (Eq. S17) evaluated on the BoltzGen refold was additionally included in selecting the top 300, with a threshold of 2.0 Å.

Each of these 300 designs were independently predicted with Protenix-v1 using ten seeds [15]. ipSAE was calculated with a predicted-aligned-error cutoff of 10 Å [16]. A self consistency check was imposed requiring a target-aligned CDR RMSD of 2 Å between the highest ipSAE Protenix-v1 refold and the backbone. For the two MIMOSA<sub>BG</sub> cohorts, the selected Protenix-v1 prediction was again required to have a template-recovery RMSD below 2.0 Å. Candidates in all three cohorts were then ranked by ipSAE, and the top 32 designs from each cohort were selected for experimental testing, yielding 32 submitted BoltzGen designs and 64 submitted MIMOSA<sub>BG</sub> designs against KEAP1.

#### A5.5 BoltzGen and MIMOSA<sub>BG</sub> IL-8 sampling, filtering, and selection

For IL-8, 100,000 designs were generated under each of three protocols: unconstrained BoltzGen, MIMOSA<sub>BG</sub> with 8IC0 and 6WZM templates. For MIMOSA<sub>BG</sub> cohorts, at each sample a random subpatch was drawn with the following protocol: pick a residue from the interaction motif at random, and a random radius ($5 < x < 10$ Å), and include all residues within that as a mimic template. This allowed for stochastically sampling various interaction patterns, being hosted with different CDR lengths. Following repository defaults, each backbone was inverse folded once and refolded. Candidates were required to pass the standard BoltzGen sequence and structure filters. For the unconstrained cohort, the top 1000 designs were retained according to the native BoltzGen composite quality ranking. For each MIMOSA<sub>BG</sub> cohort, the mimic motif recovery RMSD (Eq. S17) evaluated on the BoltzGen refold was additionally included in filtering the top 1000, with a threshold of 3.0 Å.

Each of these top 1,000 designs was then independently refolded with Protenix-v2 [17] with 50 seeds. ipSAE was calculated with a predicted-aligned-error cutoff of 10 Å, and the top 10 refolds by ipSAE were used for downstream analysis. Among those, a design was retained if its maximum ipSAE was at least 0.70, the maximum pairwise CDR RMSD among these ten refolds was at most 3.0 Å, and the maximum target-aligned CDR RMSD to both the generated backbone and the

BoltzGen refold was at most 3.0 Å. For the two MIMOSA<sub>BG</sub> cohorts, the maximum motif-recovery RMSD was additionally required to be at most 3.0 Å. After filtering, candidates were ranked by maximum ipSAE, and the top 60 designs from each cohort were selected for experimental testing.

For MIMOSA<sub>BG</sub> cohorts in initial ranking and Protenix-v2 filtering, Eq. S17 was computed against the randomly subsampled patch for the particular design. On the other hand, the whole interaction template was used for sequence fixing, such that an unguided residue which mimicked due to the generator placing it in the correct position was still fixed in residue identity and given as context to the inverse folder.

### A6. Experimental setup

VHH designs were expressed and characterized for binding through three routes. Expression and binding characterization are described separately below, and the assignment of each design cohort to these procedures is summarized in Table S3.

#### A6.1 Expression setups

##### Expression setup E1: mammalian expression and C-tag purification.

Each VHH was encoded in a mammalian expression plasmid containing the human cytomegalovirus immediate-early enhancer/promoter with intron A, a human immunoglobulin heavy-chain 5'-untranslated region, a murine immunoglobulin heavy-chain signal sequence, the VHH coding sequence, a C-terminal AviTag for *in vivo* biotinylation, a C-terminal C-tag for purification, and a bovine growth-hormone polyadenylation sequence. The plasmid backbone contained a pUC18 origin and a β-lactamase gene for ampicillin selection in *E. coli*.

Microscale transient expression was performed in Expi293 cells in two 4-mL cultures using the source laboratory's standard protocol. Cells were removed by centrifugation and the supernatants were passed through a 1.2-μm AcroPrep Advance 96 Well, 1-mL vacuum filtration plate (PALL). Purification was automated on a Tecan Freedom EVO workstation using prepacked RoboColumns (Repligen); filtered supernatants were loaded onto CaptureSelect C-tag affinity resin (Thermo Fisher Scientific). Resin was equilibrated and washed with phosphate-buffered saline comprising 10 mM sodium phosphate, 1 mM potassium phosphate, 137 mM NaCl, and 2.7 mM KCl, pH 7.4. Bound VHH was eluted with 100 mM sodium citrate, pH 2.8, and immediately neutralized with 1.5 M Tris, pH 7.5. Samples with sufficient expression underwent size-exclusion chromatography on Superdex 200 (Cytiva) in 20 mM histidine and 140 mM NaCl, pH 6.0. Purity and integrity were assessed under reducing and non-reducing conditions by capillary electrophoresis-SDS on a LabChip GXII using a LabChip HT Protein Express Chip (Revvity) and the standard protein-analysis protocol.

##### Expression setup E2: cell-free expression from pre-assembled genes.

VHH amino-acid sequences were reverse-translated and codon-optimized for manufacturability, yield, and expression efficiency. The pre-assembled gene constructs carrying a C-terminal assay tag were ordered from Twist Bioscience. The constructs used in Binding setup B2 carried a C-terminal Twin-Strep tag.

Cell-free expression used an optimized prokaryotic *in vitro* translation system in 8-μL reactions containing 4 nM pre-assembled gene fragment. Reactions were incubated for 8 h at 37°C. After expression, protein concentration and yield were quantified using an affinity-based quantification assay.

##### Expression setup E3: *E. coli* expression and His-tag affinity purification.

VHH amino-acid sequences were reverse-translated and codon-optimized for bacterial expression. The VHHs were ordered as clonal genes from Twist Biosciences in a bacterial expression vector carrying C-terminal V5, c-Myc, and His tags. Expression was performed in *E. coli* in 20-mL TB-medium cultures using the source laboratory's standard protocol. *E. coli* cells were removed by centrifugation, and the VHHs were purified from the supernatant using His-tag affinity chromatography. The resin was equilibrated and washed with phosphate-buffered saline comprising 50 mM NaH<sub>2</sub>PO<sub>4</sub>,

300 mM NaCl, and 10 mM imidazole, pH 8.0. After loading the supernatant, nonspecifically bound proteins were removed by washing the resin with 1 M NaCl in 10 mM Tris, pH 6.8. Bound VHHs were eluted with 100 mM glycine, pH 2.0, and immediately neutralized with 1 M Tris, pH 8.0.

### A6.2 Binding-characterization setups

#### Binding setup B1: biotin capture and Carterra Ultra SPR.

Binding was measured on a Carterra Ultra surface-plasmon-resonance (SPR) instrument operated with Navigator Software v2.5.0.929. HBS-EP+ running buffer contained 10 mM HEPES, 150 mM NaCl, 3 mM EDTA, and 0.05% (w/v) surfactant P20, pH 7.4 (Cytiva BR100669). Biotinylated VHHs were captured as ligands with Carterra RSA200M sensor-chip kit no. 4365. The surface was functionalized by a 20-min single-channel injection of 300 nM RSA reagent. VHHs were printed with the multichannel head under bidirectional flow for 3 min for the uPA cohort and 10 min for the KEAP1 cohort.

Recombinant human uPA (R&D Systems, 1310-SE) or recombinant human KEAP1 (Sino Biological, 11981-H20B) was injected as analyte through the single-channel fluidic path at 25°C in a single-cycle kinetics format. Three increasing concentrations (33.3, 100, and 300 nM) were injected for 3 min each, with a 30-min dissociation after each injection. Between complete cycles, the surface was regenerated through the single-channel fluidic path with two consecutive 1-min injections of 6 M guanidine hydrochloride in 0.25 M NaOH. Sensorgrams were double-referenced against a buffer blank and several reference spots and analyzed with Carterra Kinetics Evaluation Software v2.0.0.5078 and GraphPad Prism v10.5.0. The supplied assay methodology does not specify a kinetic model or numerical binder-call threshold. In the final experimental source tables, MIMOSA<sub>RFA</sub> designs were classified from the normalized on-target capture level using a threshold of 0.2. No fitted kinetic parameters were reported for these cohorts.

#### Binding setup B2: Twin-Strep capture and Carterra LSA XT SPR.

Binding was measured by SPR on a Carterra LSA XT. A carboxymethylated sensor chip was conditioned with 50 mM NaOH and activated with freshly prepared 1-ethyl-3-(3-dimethylaminopropyl)carbodiimide/*N*-hydroxysulfosuccinimide (EDC/NHS; Xantec). Strep-Tactin XT (50 µg/mL; IBA Lifesciences) in 10 mM sodium acetate, pH 4.5, was covalently coupled to the reference and capture sensors. Remaining reactive groups were quenched with 1 M ethanolamine hydrochloride, approximately pH 8.5. For these experiments, the post-immobilization wash comprised three 1-min washes with 10 mM glycine-HCl. Reference and capture sensors underwent identical treatment.

Twin-Strep-tagged VHHs were captured with the 96-channel multichannel head under bidirectional flow for 750 s, followed by a 600-s baseline in running buffer. The running buffer was 10 mM HEPES, 150 mM NaCl, 3 mM EDTA, and 0.05% Tween-20, pH 7.4, and the flow rate was 50 µL/min. The supplied methodology describes a seven-concentration, half-log dilution series. At each increasing concentration, the assay comprised a 60-s buffer baseline, 300-s association, and 600-s dissociation. The concentrations were applied as a single-cycle series without intermediate regeneration.

After each complete concentration series, the surface was regenerated with two 30-s injections of 10 mM glycine-HCl, pH 1.5, followed by two 30-s injections of 3 M guanidine hydrochloride and a final 5-min wash in running buffer.

Sensorgrams were processed with Adaptyv Fitting software. Traces were trimmed to the association, dissociation, and baseline phases, corrected for transition jumps, aligned, and baseline- and reference-subtracted. The methodology specifies global fitting to a 1:1 Langmuir model, with $k_{\text{off}}$ and $K_D$ fitted directly and $k_{\text{on}} = k_{\text{off}}/K_D$. Fits were attempted in a prioritized order using global, full, dissociation-only, or slope-based individual fits, followed when necessary by group-level equilibrium/saturation, constant/flat, or semilog-linear models. Fits were scored and filtered with quality metrics, and the final kinetic parameters were selected according to fit quality. A design was called a binder when it produced quantifiable binding curves and a calculated $K_D$. When reliable fitting was not possible, an association-phase signal shift of at least 300% over the negative control could support a binding call.

Although the methodology identifies 1:1 Langmuir fitting as the nominal global analysis, the selected reporting model in the campaign export was a bivalent-analyte model for 23 of the 24 final IL-8 binders and a conformational model for the remaining binder. Those IL-8 values are therefore reported as apparent affinities, $K_{D,\text{app}}$, rather than intrinsic monovalent affinities.

#### Binding setup B3: anti-V5 capture and 1:1 SPR analysis.

Binding was measured at 25°C on a Carterra LSA XT surface-plasmon-resonance (SPR) instrument operated with Navigator Software v2.1.2.3429. The VHHs were captured for 12 min through their C-terminal V5 tag on an HC30M sensor chip functionalized with a mouse IgG1 anti-V5 antibody (Chromotek, #SV5-PK1). Recombinant human IL-8 was injected as the analyte in a single-cycle kinetics format over an increasing concentration series in HBS-EP+ running buffer. Five increasing concentrations (0, 18.5, 56.6, 167, and 500 nM) were injected for 3 min each, with a 5-min dissociation after each injection. Between complete cycles, the surface was regenerated through the single-channel fluidic path with three consecutive 1-min injections of 10 mM glycine-HCl, pH 1.9. Sensorgrams were double-referenced and globally fitted to a 1:1 Langmuir binding model with Carterra Kinetics Evaluation Software v2.0.05078.

### A6.3 Assignment of experimental setups to design cohorts

Table S3: Mapping of computational cohorts to the expression and binding-characterization setups defined above.

| Design cohort | Expression setup | Binding setup |
| --- | --- | --- |
| MIMOSA<sub>RFA</sub> KEAP1 and uPA | E1 | B1 |
| BoltzGen and MIMOSA<sub>BG</sub> KEAP1 | E2 | B2 |
| MIMOSA<sub>BG</sub> IL-8 | E2 | B2 |
| BoltzGen IL-8 | E3 | B3 |

### A7. Post-hoc sequence and cohort analysis

The post-hoc population comprises the 395 designs submitted for experimental testing. Four IL-8 MIMOSA<sub>BG</sub> constructs that could not be loaded onto the sensor are retained in submitted-cohort summaries but excluded from binder-versus-non-binder comparisons, leaving 391 assayable designs.

Sequences were numbered and CDR regions are defined with ANARCI v2.0.8 using the IMGT numbering scheme and `mode=accuracy`. Within-cohort sequence diversity is quantified as mean pairwise normalized Levenshtein distance. For sequence pair $a, b$, loop-specific distance is $L(a, b) / \max(|a|, |b|)$. Combined-CDR distance is the sum of the three loop edit counts divided by the sum of their pairwise maximum lengths.

### B. Supplementary Results

#### B1. Composition of the experimental cohorts

Overall, the submitted set of designs contained 395 VHHs: 180 targeting IL-8, 158 targeting KEAP1, and 57 targeting uPA (Table S4). Of these, 391 were assayable and yielded 68 binders and 323 non-binders. Four IL-8 MIMOSA<sub>BG</sub> constructs could not be loaded onto the sensor. These four are counted in the submitted-cohort denominators for hit-rate calculations and in the sequence summaries, but are reported separately from assayed non-binders.

Table S4: Composition and experimental outcomes of the submitted cohorts. The Template column identifies the interaction template used for mimicry. Percentages in the binder column use all submitted designs as the denominator.

| Target | Workflow | Condition | Template | Submitted | Assayable | Binders | Non-binders | Not assayable |
| --- | --- | --- | --- | --- | --- | --- | --- | --- |
| uPA | MIMOSA<sub>RFA</sub> | Geometry-only | 3PB1 | 18 | 18 | 7 (38.9%) | 11 | 0 |
| uPA | MIMOSA<sub>RFA</sub> | Sequence-fixed | 3PB1 | 39 | 39 | 8 (20.5%) | 31 | 0 |
| KEAP1 | MIMOSA<sub>RFA</sub> | Geometry-only | 2FLU | 38 | 38 | 4 (10.5%) | 34 | 0 |
| KEAP1 | MIMOSA<sub>RFA</sub> | Sequence-fixed | 2FLU | 24 | 24 | 11 (45.8%) | 13 | 0 |
| KEAP1 | MIMOSA<sub>BG</sub> | Geometry-only | 2FLU | 32 | 32 | 2 (6.3%) | 30 | 0 |
| KEAP1 | MIMOSA<sub>BG</sub> | Sequence-fixed | 2FLU | 32 | 32 | 12 (37.5%) | 20 | 0 |
| IL-8 | MIMOSA<sub>BG</sub> | Sequence-fixed | 6WZM | 60 | 58 | 12 (20.0%) | 46 | 2 |
| IL-8 | MIMOSA<sub>BG</sub> | Sequence-fixed | 8IC0 | 60 | 58 | 12 (20.0%) | 46 | 2 |
| KEAP1 | BoltzGen | Unconstrained | N/A | 32 | 32 | 0 (0.0%) | 32 | 0 |
| IL-8 | BoltzGen | Unconstrained | N/A | 60 | 60 | 0 (0.0%) | 60 | 0 |
| Total (MIMOSA cohorts only) |  |  |  | 303 | 299 | 68 (22.4%) | 231 | 4 |
| Total |  |  |  | 395 | 391 | 68 (17.2%) | 323 | 4 |

#### B2. Framework and CDR-length composition of the submitted cohorts

Across all submissions, IMGT CDR-H1 ranged from 5 to 11 residues, CDR-H2 from 4 to 10, and CDR-H3 from 6 to 27. As described in Section A5.1, the BoltzGen and MIMOSA<sub>BG</sub> campaigns drew from a five-framework menu and varied all three loop lengths, whereas the MIMOSA<sub>RFA</sub> campaigns used only framework 3eak, fixing H1 and H2 at length 8 and varying only H3 length. Accordingly, loop-length coverage should be interpreted within each workflow and campaign, not as a direct comparison between MIMOSA<sub>BG</sub> and MIMOSA<sub>RFA</sub>. Within their respective campaigns, the submitted MIMOSA cohorts retained broad loop-length coverage (Table S5). The 120 IL-8 MIMOSA<sub>BG</sub> designs occupied 82 distinct H1/H2/H3 length triples across three of the five available frameworks, without any a priori framework restriction. The 24 binders spanned 18 triples across all three retained frameworks. The 64 KEAP1 MIMOSA<sub>BG</sub> designs occupied 52 triples, and all 14 binders had different triples, spanning three different frameworks. Because MIMOSA<sub>RFA</sub> fixed H1 and H2 and varied only H3, its submitted cohorts covered a narrower set of loop architectures, with six H3 lengths on KEAP1 and three on uPA.

Table S5: Framework composition, IMGT CDR lengths, and the number of distinct H1/H2/H3 length triples per submitted condition. Lengths are median [minimum–maximum].

| Target | Workflow | Condition | Template | Framework counts | H1 | H2 | H3 | Triples |
| --- | --- | --- | --- | --- | --- | --- | --- | --- |
| uPA | MIMOSA<sub>RFA</sub> | Geometry-only | 3PB1 | **3eak**:18 | 8 [8–8] | 8 [8–8] | 13 [12–13] | 2 |
| uPA | MIMOSA<sub>RFA</sub> | Sequence-fixed | 3PB1 | **3eak**:39 | 8 [8–8] | 8 [8–8] | 13 [11–13] | 3 |
| KEAP1 | MIMOSA<sub>RFA</sub> | Geometry-only | 2FLU | **3eak**:38 | 8 [8–8] | 8 [8–8] | 18 [10–19] | 6 |
| KEAP1 | MIMOSA<sub>RFA</sub> | Sequence-fixed | 2FLU | **3eak**:24 | 8 [8–8] | 8 [8–8] | 18 [10–19] | 4 |
| KEAP1 | MIMOSA<sub>BG</sub> | Geometry-only | 2FLU | **7eow**:17; **7x10**:3; **8coh**:9; **8z8v**:3 | 8 [6–10] | 8 [6–10] | 21 [8–27] | 26 |
| KEAP1 | MIMOSA<sub>BG</sub> | Sequence-fixed | 2FLU | **7eow**:21; **7x10**:2; **8coh**:9 | 8 [6–10] | 7 [5–10] | 22 [12–26] | 32 |
| IL-8 | MIMOSA<sub>BG</sub> | Sequence-fixed | 6WZM | **3eak**:29; **8coh**:18; **8z8v**:13 | 9 [6–11] | 8 [6–10] | 20 [11–26] | 44 |
| IL-8 | MIMOSA<sub>BG</sub> | Sequence-fixed | 8IC0 | **3eak**:12; **8coh**:38; **8z8v**:10 | 8 [6–11] | 8 [6–10] | 20 [6–24] | 44 |
| KEAP1 | BoltzGen | Unconstrained | N/A | **7eow**:11; **7x10**:11; **8coh**:3; **8z8v**:7 | 8 [6–10] | 8 [5–10] | 20 [8–25] | 30 |
| IL-8 | BoltzGen | Unconstrained | N/A | **7eow**:47; **8coh**:11; **7x10**:1; **8z8v**:1 | 5 [5–9] | 6 [4–9] | 19 [9–27] | 27 |

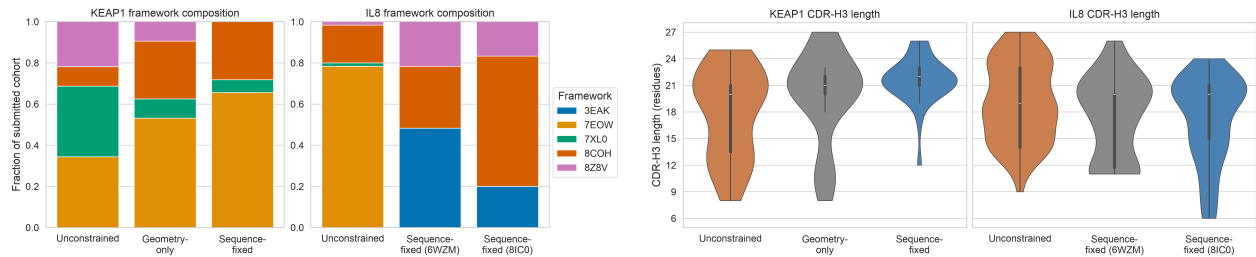

(a) Framework composition of the submitted Boltz-Gen and MIMOSA<sub>BG</sub> cohorts.

(b) IMGT CDR-H3 length distributions in the Boltz-Gen and MIMOSA<sub>BG</sub> cohorts.

Figure S1: **Submitted framework and CDR-H3 length distributions.** Framework and CDR-H3 length are set jointly by the scaffold configurations; the two panels should be interpreted accordingly. These distributions describe final submissions and reflect the combined effects of generation, filtering, and selection.

#### B3. Binding outcomes by framework and CDR length

On IL-8, the MIMOSA<sub>BG</sub> cohorts contained no **7eow** designs, while the unconstrained cohort was dominated by **7eow** (47 of 60; Table S5). Different frameworks produced very different MIMOSA<sub>BG</sub> hit rates, with 1/37 for **3eak**, 14/56 for **8coh**, and 9/23 for **8z8v** among assayable designs. On the two frameworks shared by both cohorts (**8coh** and **8z8v**), MIMOSA<sub>BG</sub> produced 23/79 binders versus 0/12 for unconstrained BoltzGen. Because the framework distributions in the two cohorts emerged from filtering and selection rather than being held constant, and because framework choice is coupled to loop lengths, this comparison describes effects of the end-to-end campaign rather than an isolated effect of guidance.

All MIMOSA<sub>RFA</sub> designs used a single framework, so within-campaign outcome differences come from other factors. For KEAP1 MIMOSA<sub>RFA</sub>, 12 of 20 designs at IMGT H3 length 12 bound compared with 3 of 42 designs at all other H3 lengths. Within H3 length 12, 9/9 sequence-fixed designs bound versus 3/11 geometry-only designs.

To identify which CDRs hosted the scattered IL-8 motifs, we aligned each complete interaction template to the target in the top Protenix-v2 refold, recomputed the Hungarian assignment, and counted assigned pairs within 2 Å. All 24 binders used H3 to host the motif, with 16 using H3 alone (Table S6).

Table S6: CDR combinations hosting the assigned template residues in the top Protenix-v2 refold, for each of the 24 IL-8 MIMOSA<sub>BG</sub> binders.

| Interaction template | H3 | H2 + H3 | H1 + H3 | Total |
| --- | --- | --- | --- | --- |
| 6WZM | 7 | 5 | 0 | 12 |
| 8IC0 | 9 | 2 | 1 | 12 |
| Total | 16 | 7 | 1 | 24 |

##### B4. MIMOSA focuses CDR sequence space without sequence collapse

Within-cohort sequence diversity was quantified as the mean pairwise normalized Levenshtein distance between designs (Section A7). Lower sequence diversity corresponds to a more focused cohort, one whose designs occupy a smaller neighborhood of sequence space. For IL-8, both MIMOSA<sub>BG</sub> cohorts were more focused than the unconstrained cohort. Combined-CDR diversity was 12.8% lower for both 6WZM and 8IC0, and H3 diversity was 12.8% and 13.7% lower, respectively. On KEAP1, combined-CDR diversity was 8.0% lower for geometry-only MIMOSA<sub>BG</sub> and 20.2% lower for sequence-fixed MIMOSA<sub>BG</sub> than for unconstrained BoltzGen. The corresponding H3 differences were 3.8% and 26.4%. Direct fixed-versus-geometry comparisons showed the largest contraction in H3, with reductions of 23.4% for KEAP1 MIMOSA<sub>BG</sub>, 24.7% for KEAP1 MIMOSA<sub>RFA</sub>, and 45.4% for uPA MIMOSA<sub>RFA</sub> (Table S7; Figs. S2 and S3). These comparisons reflect the combined effects of generation, filtering, and selection, not the isolated effect of guidance.

Table S7: Sequence diversity of each submitted cohort, measured as the mean normalized pairwise Levenshtein distance between designs.

| Target | Workflow | Condition | Template | H3 diversity | Combined-CDR diversity |
| --- | --- | --- | --- | --- | --- |
| uPA | MIMOSA<sub>RFA</sub> | Geometry-only | 3PB1 | 0.523 | 0.429 |
| uPA | MIMOSA<sub>RFA</sub> | Sequence-fixed | 3PB1 | 0.285 | 0.320 |
| KEAP1 | MIMOSA<sub>RFA</sub> | Geometry-only | 2FLU | 0.596 | 0.519 |
| KEAP1 | MIMOSA<sub>RFA</sub> | Sequence-fixed | 2FLU | 0.449 | 0.452 |
| KEAP1 | MIMOSA<sub>BG</sub> | Geometry-only | 2FLU | 0.641 | 0.607 |
| KEAP1 | MIMOSA<sub>BG</sub> | Sequence-fixed | 2FLU | 0.491 | 0.526 |
| IL-8 | MIMOSA<sub>BG</sub> | Sequence-fixed | 6WZM | 0.609 | 0.600 |
| IL-8 | MIMOSA<sub>BG</sub> | Sequence-fixed | 8IC0 | 0.603 | 0.599 |
| KEAP1 | BoltzGen | Unconstrained | N/A | 0.666 | 0.659 |
| IL-8 | BoltzGen | Unconstrained | N/A | 0.698 | 0.688 |

Focusing did not amount to sequence collapse. Normalized mean pairwise combined-CDR edit distances remained between 0.32 and 0.61 across the eight MIMOSA cohorts (Table S7), and even the most focused cohort, uPA MIMOSA<sub>RFA</sub> sequence-fixed, had a normalized mean pairwise H3 edit distance of 0.29. All 395 submitted full-domain sequences and concatenated CDR repertoires were unique, 388 distinct H3 sequences were observed, and all 68 experimental binders were unique at the full-domain, concatenated-CDRs, and H3 levels. Focusing was also not a proxy for binding. uPA showed the strongest fixed-versus-geometry H3 contraction even though fixing did not improve its hit rate, and IL-8 binders were no less diverse than non-binders in combined-CDR space: mean pairwise normalized distances were 0.609 versus 0.597 for the 6WZM cohort and 0.623 versus 0.587 for the 8IC0 one, respectively. The loop-wise pattern was consistent with motif topology: focusing for the contiguous KEAP1 and uPA motifs was concentrated in the CDR-H3 hosting loop, whereas the scattered IL-8 motifs, whose residues could be assigned across loops, were associated with reductions across multiple CDRs (Fig. S3).

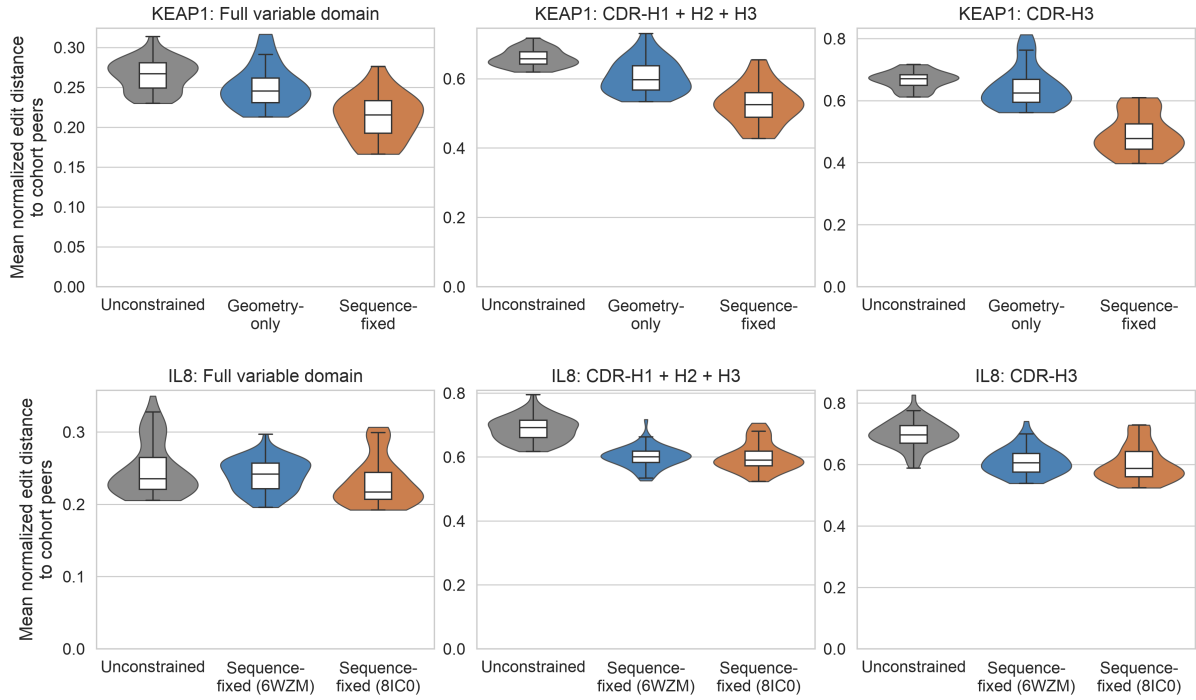

Figure S2: **Sequence diversity within the BoltzGen-generated cohorts.** For each design, we computed the mean normalized Levenshtein distance to the rest of its cohort. Violins show the distribution of these per-design distances for the full VHH, combined IMGT CDRs, and CDR-H3.

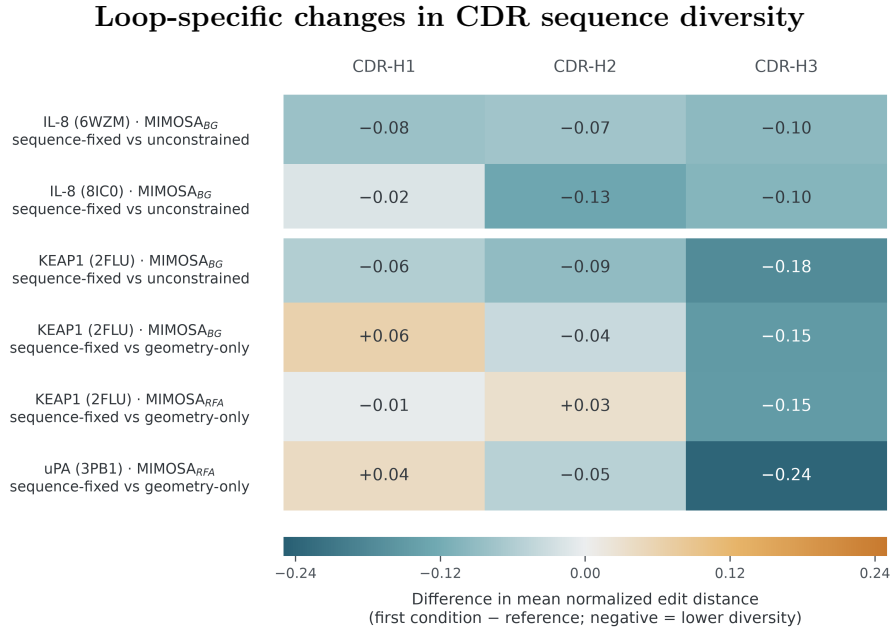

Figure S3: Each cell is the difference in mean normalized pairwise Levenshtein distance between the first condition and its reference among submitted designs. Negative values indicate a more focused repertoire. For the IL-8 cohorts and sequence-fixed KEAP1 MIMOSA<sub>BG</sub>, the reference is target-matched unconstrained BoltzGen; the remaining comparisons use the corresponding geometry-only MIMOSA cohort. These differences describe the complete generation, filtering, and selection pipelines rather than the isolated effect of guidance.

### B5. Sequence fixing has a target-dependent relationship with binding

For KEAP1, sequence fixing sharply improved binder recovery. MIMOSA<sub>BG</sub> recovered 12/32 binders with sequence fixing versus 2/32 with geometry-only guidance, while MIMOSA<sub>RFA</sub> recovered 11/24 versus 4/38. This is consistent with the chemical importance of the two acidic residues in the Nrf2 ETGE interaction. All six geometry-only KEAP1 binders across both implementations all independently recovered at least one of those two acidic identities, and three recovered both.

The pattern differed on uPA. Geometry-only MIMOSA<sub>RFA</sub> yielded 7/18 binders and sequence fixing yielded 8/39. All seven geometry-only binders independently recovered a positive residue at the interaction template’s essential position (six arginines and one lysine), despite not being required to retain the full six-residue template sequence. Together with the strong diversity contraction in the fixed uPA population, this shows that constraining more identities than strictly necessary can narrow the search without improving recovery. The choice of which mimic-motif residues to preserve is critical and depends on both the target and the source ligand.

### B6. Protenix-v2 folding confidence across cohorts

Design filtering relied on various folding models (Section A5). To place designs on a common footing, we refolded all of them with Protenix-v2 [17]. For all downstream analyses, we selected the structure with the highest ipSAE [16] across the 50 seeds. We recorded the maximum ipSAE and ipTM scores per design. Across all 395 designs, ipTM and ipSAE were strongly correlated (Pearson $r = 0.973$), so we report only the latter.

Generation methodology had a target-dependent relationship with folding confidence (Fig. S4). On KEAP1, more constrained protocols yielded higher median ipSAE. On uPA, sequence fixing lowered the median, and the three IL-8 cohorts had similar medians (Table S8).

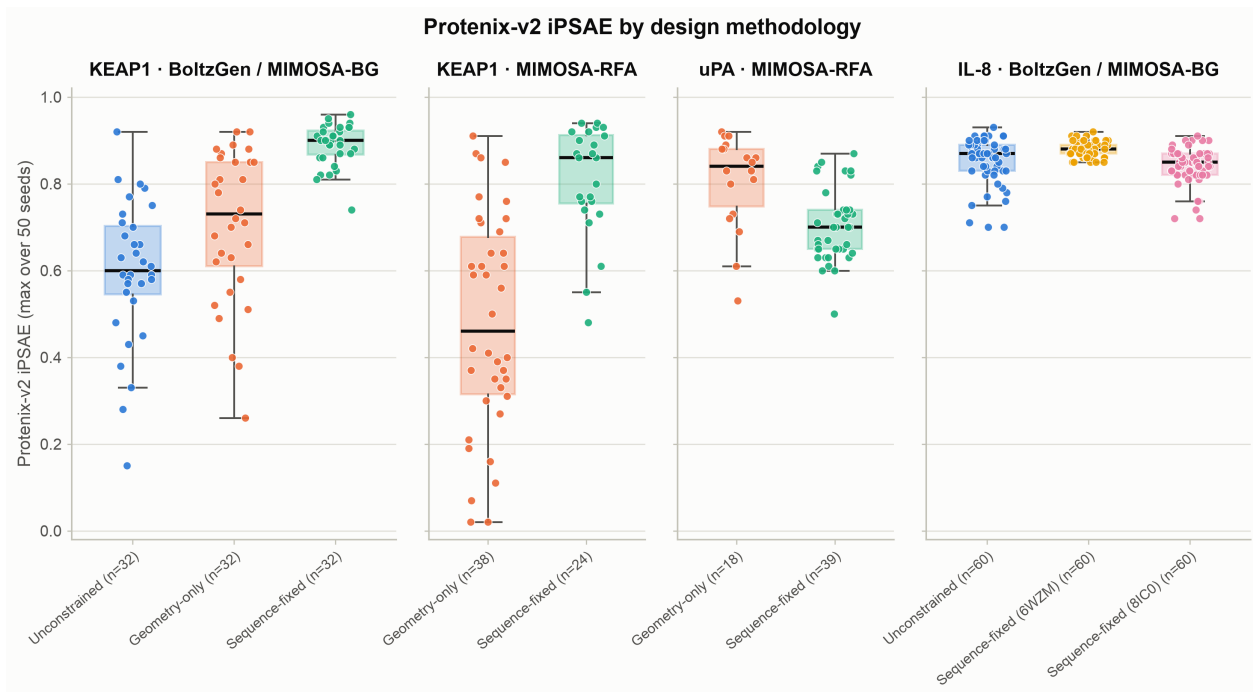

Figure S4: **Protenix-v2 ipSAE distributions by generation methodology.** The four panels show KEAP1 (BoltzGen and MIMOSA<sub>BG</sub>), KEAP1 (MIMOSA<sub>RFA</sub>: geometry-only and sequence-fixed), uPA (MIMOSA<sub>RFA</sub>: geometry-only and sequence-fixed), and IL-8 (BoltzGen and MIMOSA<sub>BG</sub> with the 6WZM and 8IC0 templates). Within each panel, cohorts are ordered by constraint level.

We also asked whether ipSAE distinguishes binders from non-binders within cohorts. It does so on KEAP1 (Pearson $r \approx 0.40$), not on uPA ($r \approx 0$) and weakly on IL-8 (Fig. S5; Table S8). Among the MIMOSA<sub>BG</sub> binders with a reported $K_D$, within-target Spearman correlations between ipSAE and $K_D$ were weak, 0.15 for IL-8 and $-0.26$ for KEAP1 (Fig. S6).

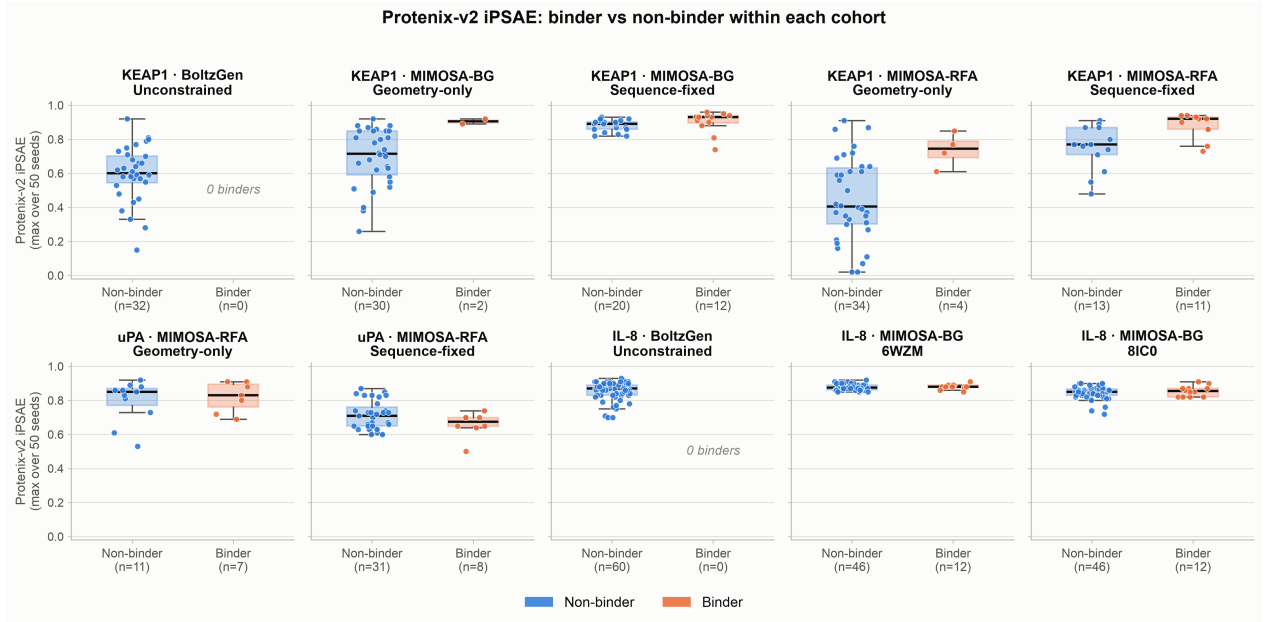

Figure S5: **Protenix-v2 ipSAE for binders versus non-binders.** Distributions are shown separately for the binders and non-binders within each of the ten submitted cohorts.

Table S8: **Protenix-v2 ipSAE score summary by cohort.** Values are the maximum ipSAE over 50 Protenix-v2 refold seeds per design. Across all 395 designs ipTM is highly collinear with ipSAE (Pearson $r = 0.973$), so only ipSAE is shown. The $n$ column is the full submitted cohort; binder and non-binder medians exclude the 4 IL-8 non-assayable constructs. Empty groups are shown as “\_”.

| Target | Workflow | Condition | Template | $n$ | Binders | Median ipSAE | Range (min–max) | Median ipSAE (binders) | Median ipSAE (non-binders) |
| --- | --- | --- | --- | --- | --- | --- | --- | --- | --- |
| uPA | MIMOSA<sub>RFA</sub> | Geometry-only | 3PB1 | 18 | 7 | 0.84 | 0.53–0.92 | 0.83 | 0.85 |
| uPA | MIMOSA<sub>RFA</sub> | Sequence-fixed | 3PB1 | 39 | 8 | 0.70 | 0.50–0.87 | 0.68 | 0.71 |
| KEAP1 | MIMOSA<sub>RFA</sub> | Geometry-only | 2FLU | 38 | 4 | 0.46 | 0.02–0.91 | 0.74 | 0.41 |
| KEAP1 | MIMOSA<sub>RFA</sub> | Sequence-fixed | 2FLU | 24 | 11 | 0.86 | 0.48–0.94 | 0.92 | 0.77 |
| KEAP1 | MIMOSA<sub>BG</sub> | Geometry-only | 2FLU | 32 | 2 | 0.73 | 0.26–0.92 | 0.91 | 0.72 |
| KEAP1 | MIMOSA<sub>BG</sub> | Sequence-fixed | 2FLU | 32 | 12 | 0.90 | 0.74–0.96 | 0.93 | 0.89 |
| IL-8 | MIMOSA<sub>BG</sub> | Sequence-fixed | 6WZM | 60 | 12 | 0.88 | 0.85–0.92 | 0.88 | 0.88 |
| IL-8 | MIMOSA<sub>BG</sub> | Sequence-fixed | 8IC0 | 60 | 12 | 0.85 | 0.72–0.91 | 0.86 | 0.85 |
| KEAP1 | BoltzGen | Unconstrained | N/A | 32 | 0 | 0.60 | 0.15–0.92 | – | 0.60 |
| IL-8 | BoltzGen | Unconstrained | N/A | 60 | 0 | 0.87 | 0.70–0.93 | – | 0.87 |

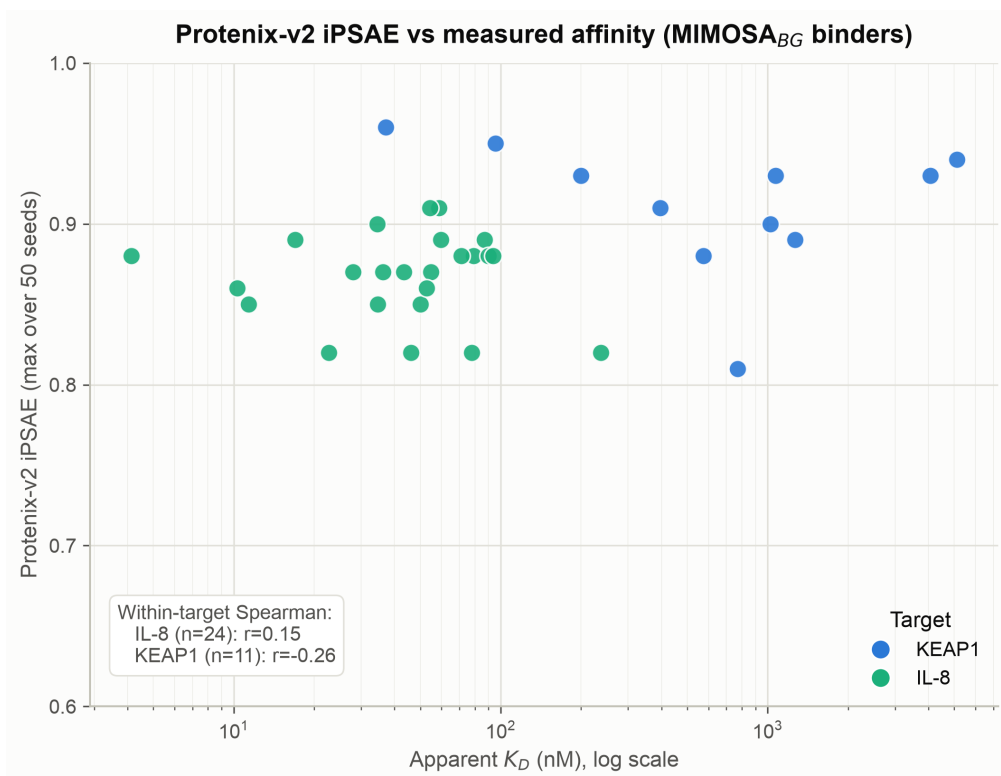

Figure S6: **Protenix-v2 ipSAE versus apparent affinity.** Per-design maximum ipSAE against apparent $K_D$ for the MIMOSA<sub>BG</sub> binders with a reported $K_D$, coloured by target; the per-target Spearman correlations are annotated on the panel.

Selection strategies differed across cohorts, which partially explains the Protenix-v2 ipSAE numbers in Table S8. The three IL-8 cohorts were selected on Protenix-v2 max-ipSAE $\geq 0.70$ (top 60 per cohort), so their ipSAE distributions are truncated below 0.70. This inflates the absolute IL-8 values and compresses the binder/non-binder separation within IL-8. The KEAP1 BoltzGen and MIMOSA<sub>BG</sub> cohorts were instead selected on Protenix-v1 ipSAE, and the MIMOSA<sub>RFA</sub> cohorts on Boltz-2 ipTM. For those cohorts, Protenix-v2 was applied only after cohort selection and lab characterization. The unified Protenix-v2 ipSAE is therefore an independent re-scoring, not the selection metric.

### B7. Closely related designs can have discordant outcomes

For each target–workflow–template cohort, we identified the nearest binder/non-binder pair (by CDR edit distance) among designs that shared strategy and framework sequence. These pairs differed by only four to six CDR edits and mimicked the same motif residues (Table S9). In the 8IC0 pair, H1 and H3 were identical and all four residue differences fell in H2. Both designs retained essentially the same transferred interaction geometry. Preserving the motif alone was therefore not sufficient to guarantee binding. Because this is an observational comparison, we cannot attribute the outcome differences to any specific residues or structural features.

Table S9: Nearest matched binder/non-binder pair in each target–workflow–template cohort. Matching required the same strategy and framework sequence. Edit counts are across concatenated IMGT CDRs.

| Target | Workflow/template | CDR edits | Locations of edits | Mimic motif retained |
| --- | --- | --- | --- | --- |
| uPA | MIMOSA<sub>RFA</sub> | 4 | Two H2 and two post-motif H3 positions | Yes |
| KEAP1 | MIMOSA<sub>RFA</sub> | 5 | Three H2 and two H3 flanking positions | Yes |
| KEAP1 | MIMOSA<sub>BG</sub> | 6 | H1 and H3 positions outside retained identities | Yes |
| IL-8 | MIMOSA<sub>BG</sub>, 6WZM | 4 | H1 and H3 positions outside retained identities | Yes |
| IL-8 | MIMOSA<sub>BG</sub>, 8IC0 | 4 | H2 only | Yes |

### B8. Sensitivity analysis for a scaffold-insertion defect in the unconstrained IL-8 cohort

At the time of our IL-8 runs, the published BoltzGen release contained two indexing defects in its scaffold-insertion parser, which we used as published. These defects were subsequently corrected upstream in commit 71cf788 of the BoltzGen repository<sup>1</sup>. We identified the defects post-hoc from the sequence and structural fingerprints of the affected designs, after experimental characterization was complete. The main-text comparison retains the full 60-design cohort to reflect the tool as published, and the sensitivity analysis below shows the findings are unchanged when restricted to the 32 canonical-cysteine designs.

The two defects operated as follows. First, a written inclusive range such as 1..5 was sampled as 0–4 because a one-based range was reused as an insertion count without the required +1. Second, H1, H2, and H3 insertions were applied sequentially, but each downstream insertion reused the original scaffold coordinate instead of adding the offsets introduced by earlier insertions.

In 28 of the 60 unconstrained IL-8 designs, these defects shifted the biological numbering of the conserved second cysteine into the CDR-H3 interval, leaving it without a canonical partner. Predicted C23–C104 Sγ–Sγ distances were 1.70–2.30 Å (median 2.11 Å) in the 32 canonical-cysteine designs, consistent with disulfide pairing, and 8.54–21.62 Å (median 16.04 Å) in the 28 affected designs. All 68 binders across the complete study had only the canonical C23 and C104 cysteines. These are observations from predicted structures. No experimental disulfide mapping was performed.

As a sensitivity check, we restricted the unconstrained IL-8 control set to the 32 designs featuring the canonical disulfide bond. All 32 remained non-binders. Against this restricted set, 24 of 116 assayable MIMOSA<sub>BG</sub> designs bound versus 0 of 32 controls, and for each interaction template the comparison was 12 of 58 versus 0 of 32. Sequence focusing was similarly preserved. Relative to the 32 canonical-cysteine controls, combined-CDR diversity was 13.7% lower for both 6WZM and 8IC0; H3 diversity was 13.6% and 14.5% lower, respectively (Table S10).

Table S10: IL-8 sensitivity analysis restricted to the 32 unconstrained designs with canonical IMGT cysteine placement and predicted C23–C104 disulfide-compatible geometry. Binder counts are shown as binders/assayable. Distances are mean normalized pairwise Levenshtein distances.

| Condition | *n* | Binders | Full VHH | H1 | H2 | H3 | Combined |
| --- | --- | --- | --- | --- | --- | --- | --- |
| Canonical control | 32 | 0/32 | 0.2720 | 0.7048 | 0.6430 | 0.7047 | 0.6948 |
| MIMOSA<sub>BG</sub>, 6WZM | 60 | 12/58 | 0.2396 | 0.6000 | 0.5691 | 0.6091 | 0.5999 |
| MIMOSA<sub>BG</sub>, 8IC0 | 60 | 12/58 | 0.2306 | 0.6647 | 0.5158 | 0.6026 | 0.5995 |

<sup>1</sup>https://github.com/HannesStark/boltzgen

### B9. Design-level summary of submitted cohorts

Tables S11, S12, S13, and S14 summarize all 395 laboratory submissions, separated by target and workflow. The tables report workflow, experimental condition, interaction template, framework, sequence-fixing status, IMGT CDR lengths, outcome, and the relevant experimental metric. For BoltzGen and MIMOSA<sub>BG</sub> designs, apparent $K_D$ values in nM are shown where reported. For MIMOSA<sub>RFA</sub> designs, normalized capture levels are reported to two decimal places for binders, while all non-binders are shown as $< 0.2$, the threshold used for binding classification. Four IL-8 designs are explicitly labeled *Not assayable* rather than merged with assayed non-binders.

Table S11: Design-level characteristics and experimental outcomes for the 57 submitted uPA MIMOSA<sub>RFA</sub> designs. Design numbers are local to this table; CDR lengths use IMGT numbering. Normalized capture levels are reported for binders; non-binders are shown as $< 0.2$, the classification threshold.

| No. | Workflow | Condition | Template | Framework | Fixed | H1 | H2 | H3 | Outcome | Normalized capture level |
| --- | --- | --- | --- | --- | --- | --- | --- | --- | --- | --- |
| 1 | MIMOSA<sub>RFA</sub> | Geometry-only | 3PB1 | 3eak | No | 8 | 8 | 13 | Binder | 1.98 |
| 2 | MIMOSA<sub>RFA</sub> | Geometry-only | 3PB1 | 3eak | No | 8 | 8 | 13 | Binder | 1.66 |
| 3 | MIMOSA<sub>RFA</sub> | Geometry-only | 3PB1 | 3eak | No | 8 | 8 | 13 | Binder | 1.62 |
| 4 | MIMOSA<sub>RFA</sub> | Geometry-only | 3PB1 | 3eak | No | 8 | 8 | 13 | Binder | 1.30 |
| 5 | MIMOSA<sub>RFA</sub> | Geometry-only | 3PB1 | 3eak | No | 8 | 8 | 13 | Binder | 0.89 |
| 6 | MIMOSA<sub>RFA</sub> | Geometry-only | 3PB1 | 3eak | No | 8 | 8 | 12 | Binder | 0.69 |
| 7 | MIMOSA<sub>RFA</sub> | Geometry-only | 3PB1 | 3eak | No | 8 | 8 | 12 | Binder | 0.29 |
| 8 | MIMOSA<sub>RFA</sub> | Geometry-only | 3PB1 | 3eak | No | 8 | 8 | 12 | Non-binder | $< 0.2$ |
| 9 | MIMOSA<sub>RFA</sub> | Geometry-only | 3PB1 | 3eak | No | 8 | 8 | 13 | Non-binder | $< 0.2$ |
| 10 | MIMOSA<sub>RFA</sub> | Geometry-only | 3PB1 | 3eak | No | 8 | 8 | 13 | Non-binder | $< 0.2$ |
| 11 | MIMOSA<sub>RFA</sub> | Geometry-only | 3PB1 | 3eak | No | 8 | 8 | 13 | Non-binder | $< 0.2$ |
| 12 | MIMOSA<sub>RFA</sub> | Geometry-only | 3PB1 | 3eak | No | 8 | 8 | 12 | Non-binder | $< 0.2$ |
| 13 | MIMOSA<sub>RFA</sub> | Geometry-only | 3PB1 | 3eak | No | 8 | 8 | 12 | Non-binder | $< 0.2$ |
| 14 | MIMOSA<sub>RFA</sub> | Geometry-only | 3PB1 | 3eak | No | 8 | 8 | 12 | Non-binder | $< 0.2$ |
| 15 | MIMOSA<sub>RFA</sub> | Geometry-only | 3PB1 | 3eak | No | 8 | 8 | 12 | Non-binder | $< 0.2$ |
| 16 | MIMOSA<sub>RFA</sub> | Geometry-only | 3PB1 | 3eak | No | 8 | 8 | 13 | Non-binder | $< 0.2$ |
| 17 | MIMOSA<sub>RFA</sub> | Geometry-only | 3PB1 | 3eak | No | 8 | 8 | 12 | Non-binder | $< 0.2$ |
| 18 | MIMOSA<sub>RFA</sub> | Geometry-only | 3PB1 | 3eak | No | 8 | 8 | 13 | Non-binder | $< 0.2$ |
| 19 | MIMOSA<sub>RFA</sub> | Sequence-fixed | 3PB1 | 3eak | Yes | 8 | 8 | 13 | Binder | 2.24 |
| 20 | MIMOSA<sub>RFA</sub> | Sequence-fixed | 3PB1 | 3eak | Yes | 8 | 8 | 13 | Binder | 1.45 |
| 21 | MIMOSA<sub>RFA</sub> | Sequence-fixed | 3PB1 | 3eak | Yes | 8 | 8 | 13 | Binder | 1.10 |
| 22 | MIMOSA<sub>RFA</sub> | Sequence-fixed | 3PB1 | 3eak | Yes | 8 | 8 | 13 | Binder | 0.96 |
| 23 | MIMOSA<sub>RFA</sub> | Sequence-fixed | 3PB1 | 3eak | Yes | 8 | 8 | 13 | Binder | 0.79 |
| 24 | MIMOSA<sub>RFA</sub> | Sequence-fixed | 3PB1 | 3eak | Yes | 8 | 8 | 13 | Binder | 0.40 |
| 25 | MIMOSA<sub>RFA</sub> | Sequence-fixed | 3PB1 | 3eak | Yes | 8 | 8 | 13 | Binder | 0.27 |
| 26 | MIMOSA<sub>RFA</sub> | Sequence-fixed | 3PB1 | 3eak | Yes | 8 | 8 | 13 | Binder | 0.21 |
| 27 | MIMOSA<sub>RFA</sub> | Sequence-fixed | 3PB1 | 3eak | Yes | 8 | 8 | 13 | Non-binder | $< 0.2$ |
| 28 | MIMOSA<sub>RFA</sub> | Sequence-fixed | 3PB1 | 3eak | Yes | 8 | 8 | 13 | Non-binder | $< 0.2$ |
| 29 | MIMOSA<sub>RFA</sub> | Sequence-fixed | 3PB1 | 3eak | Yes | 8 | 8 | 13 | Non-binder | $< 0.2$ |
| 30 | MIMOSA<sub>RFA</sub> | Sequence-fixed | 3PB1 | 3eak | Yes | 8 | 8 | 13 | Non-binder | $< 0.2$ |
| 31 | MIMOSA<sub>RFA</sub> | Sequence-fixed | 3PB1 | 3eak | Yes | 8 | 8 | 13 | Non-binder | $< 0.2$ |
| 32 | MIMOSA<sub>RFA</sub> | Sequence-fixed | 3PB1 | 3eak | Yes | 8 | 8 | 13 | Non-binder | $< 0.2$ |
| 33 | MIMOSA<sub>RFA</sub> | Sequence-fixed | 3PB1 | 3eak | Yes | 8 | 8 | 13 | Non-binder | $< 0.2$ |
| 34 | MIMOSA<sub>RFA</sub> | Sequence-fixed | 3PB1 | 3eak | Yes | 8 | 8 | 13 | Non-binder | $< 0.2$ |
| 35 | MIMOSA<sub>RFA</sub> | Sequence-fixed | 3PB1 | 3eak | Yes | 8 | 8 | 13 | Non-binder | $< 0.2$ |
| 36 | MIMOSA<sub>RFA</sub> | Sequence-fixed | 3PB1 | 3eak | Yes | 8 | 8 | 12 | Non-binder | $< 0.2$ |
| 37 | MIMOSA<sub>RFA</sub> | Sequence-fixed | 3PB1 | 3eak | Yes | 8 | 8 | 12 | Non-binder | $< 0.2$ |
| 38 | MIMOSA<sub>RFA</sub> | Sequence-fixed | 3PB1 | 3eak | Yes | 8 | 8 | 13 | Non-binder | $< 0.2$ |
| 39 | MIMOSA<sub>RFA</sub> | Sequence-fixed | 3PB1 | 3eak | Yes | 8 | 8 | 13 | Non-binder | $< 0.2$ |
| 40 | MIMOSA<sub>RFA</sub> | Sequence-fixed | 3PB1 | 3eak | Yes | 8 | 8 | 13 | Non-binder | $< 0.2$ |
| 41 | MIMOSA<sub>RFA</sub> | Sequence-fixed | 3PB1 | 3eak | Yes | 8 | 8 | 12 | Non-binder | $< 0.2$ |
| 42 | MIMOSA<sub>RFA</sub> | Sequence-fixed | 3PB1 | 3eak | Yes | 8 | 8 | 13 | Non-binder | $< 0.2$ |
| 43 | MIMOSA<sub>RFA</sub> | Sequence-fixed | 3PB1 | 3eak | Yes | 8 | 8 | 13 | Non-binder | $< 0.2$ |
| 44 | MIMOSA<sub>RFA</sub> | Sequence-fixed | 3PB1 | 3eak | Yes | 8 | 8 | 13 | Non-binder | $< 0.2$ |
| 45 | MIMOSA<sub>RFA</sub> | Sequence-fixed | 3PB1 | 3eak | Yes | 8 | 8 | 13 | Non-binder | $< 0.2$ |
| 46 | MIMOSA<sub>RFA</sub> | Sequence-fixed | 3PB1 | 3eak | Yes | 8 | 8 | 13 | Non-binder | $< 0.2$ |
| 47 | MIMOSA<sub>RFA</sub> | Sequence-fixed | 3PB1 | 3eak | Yes | 8 | 8 | 12 | Non-binder | $< 0.2$ |
| 48 | MIMOSA<sub>RFA</sub> | Sequence-fixed | 3PB1 | 3eak | Yes | 8 | 8 | 13 | Non-binder | $< 0.2$ |

Continued on next page

Table S11 (continued)

| No. | Workflow | Condition | Template | Framework | Fixed | H1 | H2 | H3 | Outcome | Normalized capture level |
| --- | --- | --- | --- | --- | --- | --- | --- | --- | --- | --- |
| 49 | MIMOSA<sub>RFA</sub> | Sequence-fixed | 3PB1 | 3eak | Yes | 8 | 8 | 13 | Non-binder | < 0.2 |
| 50 | MIMOSA<sub>RFA</sub> | Sequence-fixed | 3PB1 | 3eak | Yes | 8 | 8 | 12 | Non-binder | < 0.2 |
| 51 | MIMOSA<sub>RFA</sub> | Sequence-fixed | 3PB1 | 3eak | Yes | 8 | 8 | 13 | Non-binder | < 0.2 |
| 52 | MIMOSA<sub>RFA</sub> | Sequence-fixed | 3PB1 | 3eak | Yes | 8 | 8 | 12 | Non-binder | < 0.2 |
| 53 | MIMOSA<sub>RFA</sub> | Sequence-fixed | 3PB1 | 3eak | Yes | 8 | 8 | 11 | Non-binder | < 0.2 |
| 54 | MIMOSA<sub>RFA</sub> | Sequence-fixed | 3PB1 | 3eak | Yes | 8 | 8 | 13 | Non-binder | < 0.2 |
| 55 | MIMOSA<sub>RFA</sub> | Sequence-fixed | 3PB1 | 3eak | Yes | 8 | 8 | 13 | Non-binder | < 0.2 |
| 56 | MIMOSA<sub>RFA</sub> | Sequence-fixed | 3PB1 | 3eak | Yes | 8 | 8 | 13 | Non-binder | < 0.2 |
| 57 | MIMOSA<sub>RFA</sub> | Sequence-fixed | 3PB1 | 3eak | Yes | 8 | 8 | 13 | Non-binder | < 0.2 |

Table S12: Design-level characteristics and experimental outcomes for the 62 submitted KEAP1 MIMOSA<sub>RFA</sub> designs. Design numbers are local to this table; CDR lengths use IMGT numbering. Normalized capture levels are reported for binders; non-binders are shown as < 0.2, the classification threshold.

| No. | Workflow | Condition | Template | Framework | Fixed | H1 | H2 | H3 | Outcome | Normalized capture level |
| --- | --- | --- | --- | --- | --- | --- | --- | --- | --- | --- |
| 1 | MIMOSA<sub>RFA</sub> | Geometry-only | 2FLU | 3eak | No | 8 | 8 | 12 | Binder | 2.59 |
| 2 | MIMOSA<sub>RFA</sub> | Geometry-only | 2FLU | 3eak | No | 8 | 8 | 12 | Binder | 0.79 |
| 3 | MIMOSA<sub>RFA</sub> | Geometry-only | 2FLU | 3eak | No | 8 | 8 | 10 | Binder | 0.26 |
| 4 | MIMOSA<sub>RFA</sub> | Geometry-only | 2FLU | 3eak | No | 8 | 8 | 12 | Binder | 0.24 |
| 5 | MIMOSA<sub>RFA</sub> | Geometry-only | 2FLU | 3eak | No | 8 | 8 | 12 | Non-binder | < 0.2 |
| 6 | MIMOSA<sub>RFA</sub> | Geometry-only | 2FLU | 3eak | No | 8 | 8 | 19 | Non-binder | < 0.2 |
| 7 | MIMOSA<sub>RFA</sub> | Geometry-only | 2FLU | 3eak | No | 8 | 8 | 12 | Non-binder | < 0.2 |
| 8 | MIMOSA<sub>RFA</sub> | Geometry-only | 2FLU | 3eak | No | 8 | 8 | 10 | Non-binder | < 0.2 |
| 9 | MIMOSA<sub>RFA</sub> | Geometry-only | 2FLU | 3eak | No | 8 | 8 | 12 | Non-binder | < 0.2 |
| 10 | MIMOSA<sub>RFA</sub> | Geometry-only | 2FLU | 3eak | No | 8 | 8 | 19 | Non-binder | < 0.2 |
| 11 | MIMOSA<sub>RFA</sub> | Geometry-only | 2FLU | 3eak | No | 8 | 8 | 14 | Non-binder | < 0.2 |
| 12 | MIMOSA<sub>RFA</sub> | Geometry-only | 2FLU | 3eak | No | 8 | 8 | 18 | Non-binder | < 0.2 |
| 13 | MIMOSA<sub>RFA</sub> | Geometry-only | 2FLU | 3eak | No | 8 | 8 | 10 | Non-binder | < 0.2 |
| 14 | MIMOSA<sub>RFA</sub> | Geometry-only | 2FLU | 3eak | No | 8 | 8 | 19 | Non-binder | < 0.2 |
| 15 | MIMOSA<sub>RFA</sub> | Geometry-only | 2FLU | 3eak | No | 8 | 8 | 12 | Non-binder | < 0.2 |
| 16 | MIMOSA<sub>RFA</sub> | Geometry-only | 2FLU | 3eak | No | 8 | 8 | 18 | Non-binder | < 0.2 |
| 17 | MIMOSA<sub>RFA</sub> | Geometry-only | 2FLU | 3eak | No | 8 | 8 | 19 | Non-binder | < 0.2 |
| 18 | MIMOSA<sub>RFA</sub> | Geometry-only | 2FLU | 3eak | No | 8 | 8 | 18 | Non-binder | < 0.2 |
| 19 | MIMOSA<sub>RFA</sub> | Geometry-only | 2FLU | 3eak | No | 8 | 8 | 18 | Non-binder | < 0.2 |
| 20 | MIMOSA<sub>RFA</sub> | Geometry-only | 2FLU | 3eak | No | 8 | 8 | 19 | Non-binder | < 0.2 |
| 21 | MIMOSA<sub>RFA</sub> | Geometry-only | 2FLU | 3eak | No | 8 | 8 | 12 | Non-binder | < 0.2 |
| 22 | MIMOSA<sub>RFA</sub> | Geometry-only | 2FLU | 3eak | No | 8 | 8 | 19 | Non-binder | < 0.2 |
| 23 | MIMOSA<sub>RFA</sub> | Geometry-only | 2FLU | 3eak | No | 8 | 8 | 18 | Non-binder | < 0.2 |
| 24 | MIMOSA<sub>RFA</sub> | Geometry-only | 2FLU | 3eak | No | 8 | 8 | 18 | Non-binder | < 0.2 |
| 25 | MIMOSA<sub>RFA</sub> | Geometry-only | 2FLU | 3eak | No | 8 | 8 | 19 | Non-binder | < 0.2 |
| 26 | MIMOSA<sub>RFA</sub> | Geometry-only | 2FLU | 3eak | No | 8 | 8 | 18 | Non-binder | < 0.2 |
| 27 | MIMOSA<sub>RFA</sub> | Geometry-only | 2FLU | 3eak | No | 8 | 8 | 13 | Non-binder | < 0.2 |
| 28 | MIMOSA<sub>RFA</sub> | Geometry-only | 2FLU | 3eak | No | 8 | 8 | 19 | Non-binder | < 0.2 |
| 29 | MIMOSA<sub>RFA</sub> | Geometry-only | 2FLU | 3eak | No | 8 | 8 | 18 | Non-binder | < 0.2 |
| 30 | MIMOSA<sub>RFA</sub> | Geometry-only | 2FLU | 3eak | No | 8 | 8 | 12 | Non-binder | < 0.2 |
| 31 | MIMOSA<sub>RFA</sub> | Geometry-only | 2FLU | 3eak | No | 8 | 8 | 12 | Non-binder | < 0.2 |
| 32 | MIMOSA<sub>RFA</sub> | Geometry-only | 2FLU | 3eak | No | 8 | 8 | 18 | Non-binder | < 0.2 |
| 33 | MIMOSA<sub>RFA</sub> | Geometry-only | 2FLU | 3eak | No | 8 | 8 | 18 | Non-binder | < 0.2 |
| 34 | MIMOSA<sub>RFA</sub> | Geometry-only | 2FLU | 3eak | No | 8 | 8 | 18 | Non-binder | < 0.2 |
| 35 | MIMOSA<sub>RFA</sub> | Geometry-only | 2FLU | 3eak | No | 8 | 8 | 18 | Non-binder | < 0.2 |
| 36 | MIMOSA<sub>RFA</sub> | Geometry-only | 2FLU | 3eak | No | 8 | 8 | 12 | Non-binder | < 0.2 |
| 37 | MIMOSA<sub>RFA</sub> | Geometry-only | 2FLU | 3eak | No | 8 | 8 | 18 | Non-binder | < 0.2 |
| 38 | MIMOSA<sub>RFA</sub> | Geometry-only | 2FLU | 3eak | No | 8 | 8 | 14 | Non-binder | < 0.2 |
| 39 | MIMOSA<sub>RFA</sub> | Sequence-fixed | 2FLU | 3eak | Yes | 8 | 8 | 12 | Binder | 1.59 |
| 40 | MIMOSA<sub>RFA</sub> | Sequence-fixed | 2FLU | 3eak | Yes | 8 | 8 | 12 | Binder | 0.72 |
| 41 | MIMOSA<sub>RFA</sub> | Sequence-fixed | 2FLU | 3eak | Yes | 8 | 8 | 12 | Binder | 0.72 |
| 42 | MIMOSA<sub>RFA</sub> | Sequence-fixed | 2FLU | 3eak | Yes | 8 | 8 | 12 | Binder | 0.71 |
| 43 | MIMOSA<sub>RFA</sub> | Sequence-fixed | 2FLU | 3eak | Yes | 8 | 8 | 12 | Binder | 0.69 |
| 44 | MIMOSA<sub>RFA</sub> | Sequence-fixed | 2FLU | 3eak | Yes | 8 | 8 | 12 | Binder | 0.68 |
| 45 | MIMOSA<sub>RFA</sub> | Sequence-fixed | 2FLU | 3eak | Yes | 8 | 8 | 12 | Binder | 0.65 |

Continued on next page

Table S12 (continued)

| No. | Workflow | Condition | Template | Framework | Fixed | H1 | H2 | H3 | Outcome | Normalized capture level |
| --- | --- | --- | --- | --- | --- | --- | --- | --- | --- | --- |
| 46 | MIMOSA<sub>RFA</sub> | Sequence-fixed | 2FLU | 3eak | Yes | 8 | 8 | 12 | Binder | 0.51 |
| 47 | MIMOSA<sub>RFA</sub> | Sequence-fixed | 2FLU | 3eak | Yes | 8 | 8 | 19 | Binder | 0.48 |
| 48 | MIMOSA<sub>RFA</sub> | Sequence-fixed | 2FLU | 3eak | Yes | 8 | 8 | 12 | Binder | 0.41 |
| 49 | MIMOSA<sub>RFA</sub> | Sequence-fixed | 2FLU | 3eak | Yes | 8 | 8 | 18 | Binder | 0.21 |
| 50 | MIMOSA<sub>RFA</sub> | Sequence-fixed | 2FLU | 3eak | Yes | 8 | 8 | 19 | Non-binder | < 0.2 |
| 51 | MIMOSA<sub>RFA</sub> | Sequence-fixed | 2FLU | 3eak | Yes | 8 | 8 | 18 | Non-binder | < 0.2 |
| 52 | MIMOSA<sub>RFA</sub> | Sequence-fixed | 2FLU | 3eak | Yes | 8 | 8 | 10 | Non-binder | < 0.2 |
| 53 | MIMOSA<sub>RFA</sub> | Sequence-fixed | 2FLU | 3eak | Yes | 8 | 8 | 18 | Non-binder | < 0.2 |
| 54 | MIMOSA<sub>RFA</sub> | Sequence-fixed | 2FLU | 3eak | Yes | 8 | 8 | 18 | Non-binder | < 0.2 |
| 55 | MIMOSA<sub>RFA</sub> | Sequence-fixed | 2FLU | 3eak | Yes | 8 | 8 | 18 | Non-binder | < 0.2 |
| 56 | MIMOSA<sub>RFA</sub> | Sequence-fixed | 2FLU | 3eak | Yes | 8 | 8 | 18 | Non-binder | < 0.2 |
| 57 | MIMOSA<sub>RFA</sub> | Sequence-fixed | 2FLU | 3eak | Yes | 8 | 8 | 18 | Non-binder | < 0.2 |
| 58 | MIMOSA<sub>RFA</sub> | Sequence-fixed | 2FLU | 3eak | Yes | 8 | 8 | 18 | Non-binder | < 0.2 |
| 59 | MIMOSA<sub>RFA</sub> | Sequence-fixed | 2FLU | 3eak | Yes | 8 | 8 | 10 | Non-binder | < 0.2 |
| 60 | MIMOSA<sub>RFA</sub> | Sequence-fixed | 2FLU | 3eak | Yes | 8 | 8 | 18 | Non-binder | < 0.2 |
| 61 | MIMOSA<sub>RFA</sub> | Sequence-fixed | 2FLU | 3eak | Yes | 8 | 8 | 18 | Non-binder | < 0.2 |
| 62 | MIMOSA<sub>RFA</sub> | Sequence-fixed | 2FLU | 3eak | Yes | 8 | 8 | 19 | Non-binder | < 0.2 |

Table S13: Design-level characteristics and experimental outcomes for the 96 submitted KEAP1 BoltzGen and MIMOSA<sub>BG</sub> designs. Design numbers are local to this table; CDR lengths use IMGT numbering. Apparent $K_D$ values are reported in nM where available.

| No. | Workflow | Condition | Template | Framework | Fixed | H1 | H2 | H3 | Outcome | Apparent $K_D$ (nM) |
| --- | --- | --- | --- | --- | --- | --- | --- | --- | --- | --- |
| 1 | BoltzGen | Unconstrained | N/A | 7eow | — | 6 | 9 | 21 | Non-binder | — |
| 2 | BoltzGen | Unconstrained | N/A | 7eow | — | 6 | 10 | 25 | Non-binder | — |
| 3 | BoltzGen | Unconstrained | N/A | 7x10 | — | 8 | 6 | 21 | Non-binder | — |
| 4 | BoltzGen | Unconstrained | N/A | 7x10 | — | 7 | 8 | 20 | Non-binder | — |
| 5 | BoltzGen | Unconstrained | N/A | 8z8v | — | 10 | 10 | 11 | Non-binder | — |
| 6 | BoltzGen | Unconstrained | N/A | 7eow | — | 6 | 6 | 20 | Non-binder | — |
| 7 | BoltzGen | Unconstrained | N/A | 7x10 | — | 7 | 8 | 21 | Non-binder | — |
| 8 | BoltzGen | Unconstrained | N/A | 7x10 | — | 7 | 6 | 18 | Non-binder | — |
| 9 | BoltzGen | Unconstrained | N/A | 7x10 | — | 8 | 7 | 20 | Non-binder | — |
| 10 | BoltzGen | Unconstrained | N/A | 8coh | — | 6 | 7 | 18 | Non-binder | — |
| 11 | BoltzGen | Unconstrained | N/A | 7x10 | — | 8 | 6 | 14 | Non-binder | — |
| 12 | BoltzGen | Unconstrained | N/A | 8z8v | — | 10 | 10 | 12 | Non-binder | — |
| 13 | BoltzGen | Unconstrained | N/A | 7eow | — | 6 | 8 | 25 | Non-binder | — |
| 14 | BoltzGen | Unconstrained | N/A | 7eow | — | 7 | 6 | 21 | Non-binder | — |
| 15 | BoltzGen | Unconstrained | N/A | 7eow | — | 7 | 6 | 21 | Non-binder | — |
| 16 | BoltzGen | Unconstrained | N/A | 7eow | — | 7 | 10 | 24 | Non-binder | — |
| 17 | BoltzGen | Unconstrained | N/A | 8z8v | — | 10 | 10 | 8 | Non-binder | — |
| 18 | BoltzGen | Unconstrained | N/A | 8z8v | — | 7 | 10 | 12 | Non-binder | — |
| 19 | BoltzGen | Unconstrained | N/A | 8coh | — | 6 | 6 | 19 | Non-binder | — |
| 20 | BoltzGen | Unconstrained | N/A | 8z8v | — | 9 | 7 | 12 | Non-binder | — |
| 21 | BoltzGen | Unconstrained | N/A | 7x10 | — | 8 | 7 | 12 | Non-binder | — |
| 22 | BoltzGen | Unconstrained | N/A | 7x10 | — | 8 | 5 | 20 | Non-binder | — |
| 23 | BoltzGen | Unconstrained | N/A | 7eow | — | 6 | 10 | 21 | Non-binder | — |
| 24 | BoltzGen | Unconstrained | N/A | 7eow | — | 10 | 8 | 23 | Non-binder | — |
| 25 | BoltzGen | Unconstrained | N/A | 7x10 | — | 9 | 7 | 15 | Non-binder | — |
| 26 | BoltzGen | Unconstrained | N/A | 7x10 | — | 9 | 8 | 14 | Non-binder | — |
| 27 | BoltzGen | Unconstrained | N/A | 8coh | — | 10 | 10 | 22 | Non-binder | — |
| 28 | BoltzGen | Unconstrained | N/A | 7eow | — | 8 | 8 | 24 | Non-binder | — |
| 29 | BoltzGen | Unconstrained | N/A | 7x10 | — | 9 | 5 | 14 | Non-binder | — |
| 30 | BoltzGen | Unconstrained | N/A | 8z8v | — | 9 | 10 | 8 | Non-binder | — |
| 31 | BoltzGen | Unconstrained | N/A | 7eow | — | 6 | 10 | 24 | Non-binder | — |
| 32 | BoltzGen | Unconstrained | N/A | 8z8v | — | 9 | 7 | 12 | Non-binder | — |
| 33 | MIMOSA<sub>BG</sub> | Geometry-only | 2FLU | 7eow | No | 9 | 7 | 21 | Binder | 1264.7 |
| 34 | MIMOSA<sub>BG</sub> | Geometry-only | 2FLU | 8coh | No | 6 | 8 | 21 | Binder | — |
| 35 | MIMOSA<sub>BG</sub> | Geometry-only | 2FLU | 7xl0 | No | 8 | 6 | 11 | Non-binder | — |
| 36 | MIMOSA<sub>BG</sub> | Geometry-only | 2FLU | 7eow | No | 9 | 10 | 20 | Non-binder | — |
| 37 | MIMOSA<sub>BG</sub> | Geometry-only | 2FLU | 8z8v | No | 9 | 8 | 10 | Non-binder | — |
| 38 | MIMOSA<sub>BG</sub> | Geometry-only | 2FLU | 8coh | No | 6 | 7 | 22 | Non-binder | — |

Continued on next page

Table S13 (continued)

| No. | Workflow | Condition | Template | Framework | Fixed | H1 | H2 | H3 | Outcome | Apparent $K_D$<br>(nM) |
| --- | --- | --- | --- | --- | --- | --- | --- | --- | --- | --- |
| 39 | MIMOSA<sub>BG</sub> | Geometry-only | 2FLU | 7eow | No | 8 | 6 | 20 | Non-binder | — |
| 40 | MIMOSA<sub>BG</sub> | Geometry-only | 2FLU | 8coh | No | 7 | 8 | 26 | Non-binder | — |
| 41 | MIMOSA<sub>BG</sub> | Geometry-only | 2FLU | 7eow | No | 8 | 8 | 27 | Non-binder | — |
| 42 | MIMOSA<sub>BG</sub> | Geometry-only | 2FLU | 7eow | No | 7 | 10 | 26 | Non-binder | — |
| 43 | MIMOSA<sub>BG</sub> | Geometry-only | 2FLU | 7eow | No | 6 | 7 | 22 | Non-binder | — |
| 44 | MIMOSA<sub>BG</sub> | Geometry-only | 2FLU | 7eow | No | 8 | 8 | 20 | Non-binder | — |
| 45 | MIMOSA<sub>BG</sub> | Geometry-only | 2FLU | 8coh | No | 7 | 7 | 21 | Non-binder | — |
| 46 | MIMOSA<sub>BG</sub> | Geometry-only | 2FLU | 8coh | No | 7 | 7 | 21 | Non-binder | — |
| 47 | MIMOSA<sub>BG</sub> | Geometry-only | 2FLU | 7xl0 | No | 7 | 9 | 11 | Non-binder | — |
| 48 | MIMOSA<sub>BG</sub> | Geometry-only | 2FLU | 7eow | No | 10 | 10 | 21 | Non-binder | — |
| 49 | MIMOSA<sub>BG</sub> | Geometry-only | 2FLU | 7eow | No | 8 | 9 | 18 | Non-binder | — |
| 50 | MIMOSA<sub>BG</sub> | Geometry-only | 2FLU | 7eow | No | 9 | 9 | 20 | Non-binder | — |
| 51 | MIMOSA<sub>BG</sub> | Geometry-only | 2FLU | 7eow | No | 7 | 7 | 21 | Non-binder | — |
| 52 | MIMOSA<sub>BG</sub> | Geometry-only | 2FLU | 8coh | No | 8 | 7 | 22 | Non-binder | — |
| 53 | MIMOSA<sub>BG</sub> | Geometry-only | 2FLU | 8coh | No | 7 | 6 | 26 | Non-binder | — |
| 54 | MIMOSA<sub>BG</sub> | Geometry-only | 2FLU | 8coh | No | 6 | 10 | 20 | Non-binder | — |
| 55 | MIMOSA<sub>BG</sub> | Geometry-only | 2FLU | 7eow | No | 9 | 9 | 22 | Non-binder | — |
| 56 | MIMOSA<sub>BG</sub> | Geometry-only | 2FLU | 7eow | No | 6 | 6 | 23 | Non-binder | — |
| 57 | MIMOSA<sub>BG</sub> | Geometry-only | 2FLU | 7eow | No | 6 | 8 | 21 | Non-binder | — |
| 58 | MIMOSA<sub>BG</sub> | Geometry-only | 2FLU | 8coh | No | 7 | 7 | 20 | Non-binder | — |
| 59 | MIMOSA<sub>BG</sub> | Geometry-only | 2FLU | 8z8v | No | 8 | 7 | 8 | Non-binder | — |
| 60 | MIMOSA<sub>BG</sub> | Geometry-only | 2FLU | 7eow | No | 9 | 8 | 22 | Non-binder | — |
| 61 | MIMOSA<sub>BG</sub> | Geometry-only | 2FLU | 7eow | No | 8 | 8 | 23 | Non-binder | — |
| 62 | MIMOSA<sub>BG</sub> | Geometry-only | 2FLU | 7xl0 | No | 8 | 6 | 11 | Non-binder | — |
| 63 | MIMOSA<sub>BG</sub> | Geometry-only | 2FLU | 8z8v | No | 9 | 6 | 11 | Non-binder | — |
| 64 | MIMOSA<sub>BG</sub> | Geometry-only | 2FLU | 7eow | No | 8 | 8 | 27 | Non-binder | — |
| 65 | MIMOSA<sub>BG</sub> | Sequence-fixed | 2FLU | 7xl0 | Yes | 9 | 5 | 12 | Binder | 37.1 |
| 66 | MIMOSA<sub>BG</sub> | Sequence-fixed | 2FLU | 8coh | Yes | 6 | 6 | 21 | Binder | 95.7 |
| 67 | MIMOSA<sub>BG</sub> | Sequence-fixed | 2FLU | 8coh | Yes | 6 | 8 | 22 | Binder | 200.1 |
| 68 | MIMOSA<sub>BG</sub> | Sequence-fixed | 2FLU | 7eow | Yes | 9 | 8 | 21 | Binder | 395.2 |
| 69 | MIMOSA<sub>BG</sub> | Sequence-fixed | 2FLU | 7eow | Yes | 6 | 9 | 25 | Binder | 574.4 |
| 70 | MIMOSA<sub>BG</sub> | Sequence-fixed | 2FLU | 8coh | Yes | 6 | 10 | 22 | Binder | 771.3 |
| 71 | MIMOSA<sub>BG</sub> | Sequence-fixed | 2FLU | 7eow | Yes | 8 | 6 | 23 | Binder | 1022.1 |
| 72 | MIMOSA<sub>BG</sub> | Sequence-fixed | 2FLU | 7eow | Yes | 6 | 6 | 22 | Binder | 1070.1 |
| 73 | MIMOSA<sub>BG</sub> | Sequence-fixed | 2FLU | 7eow | Yes | 10 | 9 | 21 | Binder | 4073.3 |
| 74 | MIMOSA<sub>BG</sub> | Sequence-fixed | 2FLU | 7eow | Yes | 9 | 7 | 20 | Binder | 5110.2 |
| 75 | MIMOSA<sub>BG</sub> | Sequence-fixed | 2FLU | 8coh | Yes | 6 | 7 | 22 | Binder | — |
| 76 | MIMOSA<sub>BG</sub> | Sequence-fixed | 2FLU | 7eow | Yes | 8 | 9 | 20 | Binder | — |
| 77 | MIMOSA<sub>BG</sub> | Sequence-fixed | 2FLU | 8coh | Yes | 8 | 9 | 21 | Non-binder | — |
| 78 | MIMOSA<sub>BG</sub> | Sequence-fixed | 2FLU | 8coh | Yes | 7 | 8 | 21 | Non-binder | — |
| 79 | MIMOSA<sub>BG</sub> | Sequence-fixed | 2FLU | 7eow | Yes | 7 | 7 | 24 | Non-binder | — |
| 80 | MIMOSA<sub>BG</sub> | Sequence-fixed | 2FLU | 7eow | Yes | 6 | 7 | 23 | Non-binder | — |
| 81 | MIMOSA<sub>BG</sub> | Sequence-fixed | 2FLU | 7eow | Yes | 8 | 7 | 21 | Non-binder | — |
| 82 | MIMOSA<sub>BG</sub> | Sequence-fixed | 2FLU | 7eow | Yes | 10 | 9 | 26 | Non-binder | — |
| 83 | MIMOSA<sub>BG</sub> | Sequence-fixed | 2FLU | 7eow | Yes | 10 | 7 | 19 | Non-binder | — |
| 84 | MIMOSA<sub>BG</sub> | Sequence-fixed | 2FLU | 7eow | Yes | 7 | 10 | 23 | Non-binder | — |
| 85 | MIMOSA<sub>BG</sub> | Sequence-fixed | 2FLU | 8coh | Yes | 8 | 8 | 26 | Non-binder | — |
| 86 | MIMOSA<sub>BG</sub> | Sequence-fixed | 2FLU | 7eow | Yes | 10 | 6 | 23 | Non-binder | — |
| 87 | MIMOSA<sub>BG</sub> | Sequence-fixed | 2FLU | 7eow | Yes | 8 | 6 | 22 | Non-binder | — |
| 88 | MIMOSA<sub>BG</sub> | Sequence-fixed | 2FLU | 7eow | Yes | 6 | 7 | 20 | Non-binder | — |
| 89 | MIMOSA<sub>BG</sub> | Sequence-fixed | 2FLU | 8coh | Yes | 8 | 7 | 22 | Non-binder | — |
| 90 | MIMOSA<sub>BG</sub> | Sequence-fixed | 2FLU | 8coh | Yes | 7 | 7 | 21 | Non-binder | — |
| 91 | MIMOSA<sub>BG</sub> | Sequence-fixed | 2FLU | 7xl0 | Yes | 6 | 7 | 17 | Non-binder | — |
| 92 | MIMOSA<sub>BG</sub> | Sequence-fixed | 2FLU | 7eow | Yes | 7 | 7 | 20 | Non-binder | — |
| 93 | MIMOSA<sub>BG</sub> | Sequence-fixed | 2FLU | 7eow | Yes | 8 | 9 | 26 | Non-binder | — |
| 94 | MIMOSA<sub>BG</sub> | Sequence-fixed | 2FLU | 7eow | Yes | 8 | 10 | 22 | Non-binder | — |
| 95 | MIMOSA<sub>BG</sub> | Sequence-fixed | 2FLU | 7eow | Yes | 9 | 9 | 22 | Non-binder | — |
| 96 | MIMOSA<sub>BG</sub> | Sequence-fixed | 2FLU | 7eow | Yes | 9 | 7 | 21 | Non-binder | — |

Table S14: Design-level characteristics and experimental outcomes for the 180 submitted IL-8 BoltzGen and MIMOSA<sub>BG</sub> designs. Design numbers are local to this table; CDR lengths use IMGT numbering. Four MIMOSA<sub>BG</sub> constructs that could not be loaded are distinguished from assayed non-binders. Apparent $K_D$ values are reported in nM where available.

| No. | Workflow | Condition | Template | Framework | Fixed | H1 | H2 | H3 | Outcome | Apparent $K_D$<br>(nM) |
| --- | --- | --- | --- | --- | --- | --- | --- | --- | --- | --- |
| 1 | BoltzGen | Unconstrained | N/A | 7eow | — | 7 | 5 | 19 | Non-binder | — |
| 2 | BoltzGen | Unconstrained | N/A | 7eow | — | 5 | 7 | 19 | Non-binder | — |
| 3 | BoltzGen | Unconstrained | N/A | 7eow | — | 6 | 6 | 23 | Non-binder | — |
| 4 | BoltzGen | Unconstrained | N/A | 7eow | — | 7 | 5 | 19 | Non-binder | — |
| 5 | BoltzGen | Unconstrained | N/A | 7eow | — | 7 | 5 | 18 | Non-binder | — |
| 6 | BoltzGen | Unconstrained | N/A | 7eow | — | 5 | 6 | 18 | Non-binder | — |
| 7 | BoltzGen | Unconstrained | N/A | 7eow | — | 5 | 6 | 23 | Non-binder | — |
| 8 | BoltzGen | Unconstrained | N/A | 7eow | — | 5 | 6 | 22 | Non-binder | — |
| 9 | BoltzGen | Unconstrained | N/A | 7eow | — | 6 | 8 | 14 | Non-binder | — |
| 10 | BoltzGen | Unconstrained | N/A | 7eow | — | 5 | 7 | 18 | Non-binder | — |
| 11 | BoltzGen | Unconstrained | N/A | 7eow | — | 5 | 6 | 23 | Non-binder | — |
| 12 | BoltzGen | Unconstrained | N/A | 7eow | — | 5 | 6 | 23 | Non-binder | — |
| 13 | BoltzGen | Unconstrained | N/A | 7eow | — | 6 | 8 | 14 | Non-binder | — |
| 14 | BoltzGen | Unconstrained | N/A | 7eow | — | 6 | 6 | 26 | Non-binder | — |
| 15 | BoltzGen | Unconstrained | N/A | 7eow | — | 5 | 7 | 19 | Non-binder | — |
| 16 | BoltzGen | Unconstrained | N/A | 8coh | — | 5 | 6 | 24 | Non-binder | — |
| 17 | BoltzGen | Unconstrained | N/A | 7eow | — | 5 | 7 | 19 | Non-binder | — |
| 18 | BoltzGen | Unconstrained | N/A | 7eow | — | 5 | 6 | 23 | Non-binder | — |
| 19 | BoltzGen | Unconstrained | N/A | 7eow | — | 5 | 7 | 19 | Non-binder | — |
| 20 | BoltzGen | Unconstrained | N/A | 7eow | — | 6 | 6 | 17 | Non-binder | — |
| 21 | BoltzGen | Unconstrained | N/A | 8coh | — | 5 | 8 | 18 | Non-binder | — |
| 22 | BoltzGen | Unconstrained | N/A | 7eow | — | 7 | 5 | 17 | Non-binder | — |
| 23 | BoltzGen | Unconstrained | N/A | 7eow | — | 5 | 7 | 19 | Non-binder | — |
| 24 | BoltzGen | Unconstrained | N/A | 7eow | — | 6 | 8 | 14 | Non-binder | — |
| 25 | BoltzGen | Unconstrained | N/A | 7eow | — | 6 | 9 | 14 | Non-binder | — |
| 26 | BoltzGen | Unconstrained | N/A | 7eow | — | 5 | 6 | 18 | Non-binder | — |
| 27 | BoltzGen | Unconstrained | N/A | 7eow | — | 5 | 7 | 19 | Non-binder | — |
| 28 | BoltzGen | Unconstrained | N/A | 7eow | — | 6 | 6 | 17 | Non-binder | — |
| 29 | BoltzGen | Unconstrained | N/A | 7eow | — | 5 | 6 | 23 | Non-binder | — |
| 30 | BoltzGen | Unconstrained | N/A | 7x10 | — | 9 | 4 | 9 | Non-binder | — |
| 31 | BoltzGen | Unconstrained | N/A | 8coh | — | 5 | 6 | 25 | Non-binder | — |
| 32 | BoltzGen | Unconstrained | N/A | 7eow | — | 5 | 7 | 19 | Non-binder | — |
| 33 | BoltzGen | Unconstrained | N/A | 7eow | — | 6 | 6 | 16 | Non-binder | — |
| 34 | BoltzGen | Unconstrained | N/A | 7eow | — | 5 | 6 | 23 | Non-binder | — |
| 35 | BoltzGen | Unconstrained | N/A | 7eow | — | 5 | 7 | 19 | Non-binder | — |
| 36 | BoltzGen | Unconstrained | N/A | 7eow | — | 5 | 5 | 24 | Non-binder | — |
| 37 | BoltzGen | Unconstrained | N/A | 7eow | — | 6 | 8 | 14 | Non-binder | — |
| 38 | BoltzGen | Unconstrained | N/A | 7eow | — | 5 | 5 | 24 | Non-binder | — |
| 39 | BoltzGen | Unconstrained | N/A | 7eow | — | 7 | 8 | 14 | Non-binder | — |
| 40 | BoltzGen | Unconstrained | N/A | 8coh | — | 5 | 6 | 25 | Non-binder | — |
| 41 | BoltzGen | Unconstrained | N/A | 7eow | — | 5 | 5 | 24 | Non-binder | — |
| 42 | BoltzGen | Unconstrained | N/A | 7eow | — | 6 | 9 | 14 | Non-binder | — |
| 43 | BoltzGen | Unconstrained | N/A | 8z8v | — | 5 | 5 | 16 | Non-binder | — |
| 44 | BoltzGen | Unconstrained | N/A | 8coh | — | 5 | 5 | 21 | Non-binder | — |
| 45 | BoltzGen | Unconstrained | N/A | 7eow | — | 5 | 5 | 27 | Non-binder | — |
| 46 | BoltzGen | Unconstrained | N/A | 7eow | — | 7 | 8 | 14 | Non-binder | — |
| 47 | BoltzGen | Unconstrained | N/A | 8coh | — | 7 | 9 | 12 | Non-binder | — |
| 48 | BoltzGen | Unconstrained | N/A | 8coh | — | 5 | 6 | 25 | Non-binder | — |
| 49 | BoltzGen | Unconstrained | N/A | 7eow | — | 5 | 5 | 27 | Non-binder | — |
| 50 | BoltzGen | Unconstrained | N/A | 7eow | — | 7 | 9 | 14 | Non-binder | — |
| 51 | BoltzGen | Unconstrained | N/A | 8coh | — | 5 | 6 | 25 | Non-binder | — |
| 52 | BoltzGen | Unconstrained | N/A | 7eow | — | 6 | 8 | 14 | Non-binder | — |
| 53 | BoltzGen | Unconstrained | N/A | 8coh | — | 5 | 6 | 24 | Non-binder | — |
| 54 | BoltzGen | Unconstrained | N/A | 8coh | — | 5 | 5 | 21 | Non-binder | — |
| 55 | BoltzGen | Unconstrained | N/A | 7eow | — | 7 | 8 | 14 | Non-binder | — |
| 56 | BoltzGen | Unconstrained | N/A | 7eow | — | 6 | 9 | 14 | Non-binder | — |
| 57 | BoltzGen | Unconstrained | N/A | 8coh | — | 5 | 6 | 25 | Non-binder | — |
| 58 | BoltzGen | Unconstrained | N/A | 7eow | — | 6 | 8 | 14 | Non-binder | — |
| 59 | BoltzGen | Unconstrained | N/A | 7eow | — | 7 | 6 | 14 | Non-binder | — |
| 60 | BoltzGen | Unconstrained | N/A | 7eow | — | 5 | 5 | 17 | Non-binder | — |

Continued on next page

Table S14 (continued)

| No. | Workflow | Condition | Template | Framework | Fixed | H1 | H2 | H3 | Outcome | Apparent $K_D$<br>(nM) |
| --- | --- | --- | --- | --- | --- | --- | --- | --- | --- | --- |
| 61 | MIMOSA<sub>BG</sub> | Sequence-fixed | 6WZM | 8coh | Yes | 8 | 7 | 20 | Binder | 4.1 |
| 62 | MIMOSA<sub>BG</sub> | Sequence-fixed | 6WZM | 8coh | Yes | 6 | 10 | 22 | Binder | 10.3 |
| 63 | MIMOSA<sub>BG</sub> | Sequence-fixed | 6WZM | 8coh | Yes | 7 | 8 | 20 | Binder | 17.0 |
| 64 | MIMOSA<sub>BG</sub> | Sequence-fixed | 6WZM | 8z8v | Yes | 6 | 8 | 17 | Binder | 34.6 |
| 65 | MIMOSA<sub>BG</sub> | Sequence-fixed | 6WZM | 8coh | Yes | 10 | 10 | 22 | Binder | 54.7 |
| 66 | MIMOSA<sub>BG</sub> | Sequence-fixed | 6WZM | 8z8v | Yes | 9 | 9 | 16 | Binder | 58.6 |
| 67 | MIMOSA<sub>BG</sub> | Sequence-fixed | 6WZM | 8z8v | Yes | 6 | 8 | 11 | Binder | 59.7 |
| 68 | MIMOSA<sub>BG</sub> | Sequence-fixed | 6WZM | 8coh | Yes | 8 | 7 | 20 | Binder | 71.2 |
| 69 | MIMOSA<sub>BG</sub> | Sequence-fixed | 6WZM | 8coh | Yes | 6 | 8 | 23 | Binder | 79.2 |
| 70 | MIMOSA<sub>BG</sub> | Sequence-fixed | 6WZM | 3eak | Yes | 10 | 10 | 12 | Binder | 87.0 |
| 71 | MIMOSA<sub>BG</sub> | Sequence-fixed | 6WZM | 8coh | Yes | 6 | 7 | 22 | Binder | 89.9 |
| 72 | MIMOSA<sub>BG</sub> | Sequence-fixed | 6WZM | 8z8v | Yes | 9 | 7 | 16 | Binder | 93.5 |
| 73 | MIMOSA<sub>BG</sub> | Sequence-fixed | 6WZM | 8z8v | Yes | 8 | 7 | 17 | Non-binder | — |
| 74 | MIMOSA<sub>BG</sub> | Sequence-fixed | 6WZM | 3eak | Yes | 9 | 9 | 20 | Non-binder | — |
| 75 | MIMOSA<sub>BG</sub> | Sequence-fixed | 6WZM | 3eak | Yes | 9 | 10 | 20 | Non-binder | — |
| 76 | MIMOSA<sub>BG</sub> | Sequence-fixed | 6WZM | 3eak | Yes | 9 | 7 | 20 | Non-binder | — |
| 77 | MIMOSA<sub>BG</sub> | Sequence-fixed | 6WZM | 3eak | Yes | 9 | 7 | 20 | Non-binder | — |
| 78 | MIMOSA<sub>BG</sub> | Sequence-fixed | 6WZM | 8coh | Yes | 10 | 8 | 22 | Non-binder | — |
| 79 | MIMOSA<sub>BG</sub> | Sequence-fixed | 6WZM | 3eak | Yes | 9 | 9 | 20 | Non-binder | — |
| 80 | MIMOSA<sub>BG</sub> | Sequence-fixed | 6WZM | 8coh | Yes | 7 | 6 | 22 | Non-binder | — |
| 81 | MIMOSA<sub>BG</sub> | Sequence-fixed | 6WZM | 8coh | Yes | 10 | 10 | 22 | Non-binder | — |
| 82 | MIMOSA<sub>BG</sub> | Sequence-fixed | 6WZM | 8coh | Yes | 7 | 6 | 25 | Non-binder | — |
| 83 | MIMOSA<sub>BG</sub> | Sequence-fixed | 6WZM | 3eak | Yes | 10 | 9 | 20 | Non-binder | — |
| 84 | MIMOSA<sub>BG</sub> | Sequence-fixed | 6WZM | 8coh | Yes | 10 | 6 | 22 | Non-binder | — |
| 85 | MIMOSA<sub>BG</sub> | Sequence-fixed | 6WZM | 8coh | Yes | 7 | 6 | 26 | Non-binder | — |
| 86 | MIMOSA<sub>BG</sub> | Sequence-fixed | 6WZM | 3eak | Yes | 10 | 7 | 11 | Non-binder | — |
| 87 | MIMOSA<sub>BG</sub> | Sequence-fixed | 6WZM | 3eak | Yes | 8 | 8 | 11 | Non-binder | — |
| 88 | MIMOSA<sub>BG</sub> | Sequence-fixed | 6WZM | 3eak | Yes | 8 | 6 | 11 | Non-binder | — |
| 89 | MIMOSA<sub>BG</sub> | Sequence-fixed | 6WZM | 3eak | Yes | 10 | 7 | 11 | Non-binder | — |
| 90 | MIMOSA<sub>BG</sub> | Sequence-fixed | 6WZM | 3eak | Yes | 8 | 9 | 11 | Non-binder | — |
| 91 | MIMOSA<sub>BG</sub> | Sequence-fixed | 6WZM | 3eak | Yes | 11 | 10 | 12 | Non-binder | — |
| 92 | MIMOSA<sub>BG</sub> | Sequence-fixed | 6WZM | 3eak | Yes | 11 | 10 | 11 | Non-binder | — |
| 93 | MIMOSA<sub>BG</sub> | Sequence-fixed | 6WZM | 3eak | Yes | 9 | 9 | 20 | Non-binder | — |
| 94 | MIMOSA<sub>BG</sub> | Sequence-fixed | 6WZM | 3eak | Yes | 8 | 6 | 11 | Non-binder | — |
| 95 | MIMOSA<sub>BG</sub> | Sequence-fixed | 6WZM | 8z8v | Yes | 8 | 7 | 16 | Non-binder | — |
| 96 | MIMOSA<sub>BG</sub> | Sequence-fixed | 6WZM | 3eak | Yes | 8 | 7 | 11 | Non-binder | — |
| 97 | MIMOSA<sub>BG</sub> | Sequence-fixed | 6WZM | 8z8v | Yes | 7 | 7 | 16 | Non-binder | — |
| 98 | MIMOSA<sub>BG</sub> | Sequence-fixed | 6WZM | 8coh | Yes | 8 | 9 | 26 | Non-binder | — |
| 99 | MIMOSA<sub>BG</sub> | Sequence-fixed | 6WZM | 3eak | Yes | 11 | 7 | 20 | Non-binder | — |
| 100 | MIMOSA<sub>BG</sub> | Sequence-fixed | 6WZM | 3eak | Yes | 8 | 6 | 11 | Non-binder | — |
| 101 | MIMOSA<sub>BG</sub> | Sequence-fixed | 6WZM | 8coh | Yes | 10 | 8 | 22 | Non-binder | — |
| 102 | MIMOSA<sub>BG</sub> | Sequence-fixed | 6WZM | 3eak | Yes | 8 | 7 | 11 | Non-binder | — |
| 103 | MIMOSA<sub>BG</sub> | Sequence-fixed | 6WZM | 3eak | Yes | 11 | 10 | 11 | Non-binder | — |
| 104 | MIMOSA<sub>BG</sub> | Sequence-fixed | 6WZM | 3eak | Yes | 11 | 7 | 20 | Non-binder | — |
| 105 | MIMOSA<sub>BG</sub> | Sequence-fixed | 6WZM | 8z8v | Yes | 7 | 9 | 16 | Non-binder | — |
| 106 | MIMOSA<sub>BG</sub> | Sequence-fixed | 6WZM | 3eak | Yes | 10 | 9 | 18 | Non-binder | — |
| 107 | MIMOSA<sub>BG</sub> | Sequence-fixed | 6WZM | 3eak | Yes | 10 | 10 | 20 | Non-binder | — |
| 108 | MIMOSA<sub>BG</sub> | Sequence-fixed | 6WZM | 3eak | Yes | 10 | 7 | 11 | Non-binder | — |
| 109 | MIMOSA<sub>BG</sub> | Sequence-fixed | 6WZM | 8z8v | Yes | 8 | 6 | 15 | Non-binder | — |
| 110 | MIMOSA<sub>BG</sub> | Sequence-fixed | 6WZM | 8coh | Yes | 7 | 8 | 21 | Non-binder | — |
| 111 | MIMOSA<sub>BG</sub> | Sequence-fixed | 6WZM | 3eak | Yes | 11 | 10 | 20 | Non-binder | — |
| 112 | MIMOSA<sub>BG</sub> | Sequence-fixed | 6WZM | 3eak | Yes | 10 | 8 | 20 | Non-binder | — |
| 113 | MIMOSA<sub>BG</sub> | Sequence-fixed | 6WZM | 8z8v | Yes | 7 | 10 | 17 | Non-binder | — |
| 114 | MIMOSA<sub>BG</sub> | Sequence-fixed | 6WZM | 8z8v | Yes | 6 | 7 | 14 | Non-binder | — |
| 115 | MIMOSA<sub>BG</sub> | Sequence-fixed | 6WZM | 8z8v | Yes | 10 | 7 | 11 | Non-binder | — |
| 116 | MIMOSA<sub>BG</sub> | Sequence-fixed | 6WZM | 8z8v | Yes | 7 | 10 | 17 | Non-binder | — |
| 117 | MIMOSA<sub>BG</sub> | Sequence-fixed | 6WZM | 8coh | Yes | 10 | 7 | 24 | Non-binder | — |
| 118 | MIMOSA<sub>BG</sub> | Sequence-fixed | 6WZM | 8coh | Yes | 7 | 9 | 20 | Non-binder | — |
| 119 | MIMOSA<sub>BG</sub> | Sequence-fixed | 6WZM | 3eak | Yes | 11 | 7 | 11 | Not assayable | — |
| 120 | MIMOSA<sub>BG</sub> | Sequence-fixed | 6WZM | 3eak | Yes | 9 | 9 | 20 | Not assayable | — |
| 121 | MIMOSA<sub>BG</sub> | Sequence-fixed | 8IC0 | 8coh | Yes | 7 | 9 | 20 | Binder | 11.4 |
| 122 | MIMOSA<sub>BG</sub> | Sequence-fixed | 8IC0 | 8coh | Yes | 7 | 9 | 20 | Binder | 22.7 |
| 123 | MIMOSA<sub>BG</sub> | Sequence-fixed | 8IC0 | 8coh | Yes | 8 | 7 | 15 | Binder | 28.0 |
| 124 | MIMOSA<sub>BG</sub> | Sequence-fixed | 8IC0 | 8coh | Yes | 9 | 10 | 21 | Binder | 34.4 |
| 125 | MIMOSA<sub>BG</sub> | Sequence-fixed | 8IC0 | 8z8v | Yes | 10 | 10 | 9 | Binder | 36.3 |

Continued on next page

Table S14 (continued)

| No. | Workflow | Condition | Template | Framework | Fixed | H1 | H2 | H3 | Outcome | Apparent $K_D$<br>(nM) |
| --- | --- | --- | --- | --- | --- | --- | --- | --- | --- | --- |
| 126 | MIMOSA<sub>BG</sub> | Sequence-fixed | 8IC0 | 8z8v | Yes | 10 | 10 | 9 | Binder | 43.4 |
| 127 | MIMOSA<sub>BG</sub> | Sequence-fixed | 8IC0 | 8z8v | Yes | 10 | 10 | 12 | Binder | 46.2 |
| 128 | MIMOSA<sub>BG</sub> | Sequence-fixed | 8IC0 | 8coh | Yes | 9 | 6 | 15 | Binder | 50.0 |
| 129 | MIMOSA<sub>BG</sub> | Sequence-fixed | 8IC0 | 8coh | Yes | 7 | 9 | 21 | Binder | 52.8 |
| 130 | MIMOSA<sub>BG</sub> | Sequence-fixed | 8IC0 | 8z8v | Yes | 10 | 10 | 6 | Binder | 54.3 |
| 131 | MIMOSA<sub>BG</sub> | Sequence-fixed | 8IC0 | 8coh | Yes | 8 | 7 | 20 | Binder | 77.9 |
| 132 | MIMOSA<sub>BG</sub> | Sequence-fixed | 8IC0 | 8z8v | Yes | 10 | 10 | 9 | Binder | 237.6 |
| 133 | MIMOSA<sub>BG</sub> | Sequence-fixed | 8IC0 | 3eak | Yes | 10 | 6 | 15 | Non-binder | — |
| 134 | MIMOSA<sub>BG</sub> | Sequence-fixed | 8IC0 | 8coh | Yes | 7 | 10 | 20 | Non-binder | — |
| 135 | MIMOSA<sub>BG</sub> | Sequence-fixed | 8IC0 | 3eak | Yes | 8 | 8 | 13 | Non-binder | — |
| 136 | MIMOSA<sub>BG</sub> | Sequence-fixed | 8IC0 | 8coh | Yes | 7 | 9 | 20 | Non-binder | — |
| 137 | MIMOSA<sub>BG</sub> | Sequence-fixed | 8IC0 | 8coh | Yes | 6 | 6 | 22 | Non-binder | — |
| 138 | MIMOSA<sub>BG</sub> | Sequence-fixed | 8IC0 | 8z8v | Yes | 7 | 7 | 10 | Non-binder | — |
| 139 | MIMOSA<sub>BG</sub> | Sequence-fixed | 8IC0 | 8coh | Yes | 8 | 8 | 16 | Non-binder | — |
| 140 | MIMOSA<sub>BG</sub> | Sequence-fixed | 8IC0 | 8coh | Yes | 7 | 6 | 22 | Non-binder | — |
| 141 | MIMOSA<sub>BG</sub> | Sequence-fixed | 8IC0 | 8coh | Yes | 9 | 6 | 15 | Non-binder | — |
| 142 | MIMOSA<sub>BG</sub> | Sequence-fixed | 8IC0 | 8coh | Yes | 7 | 6 | 20 | Non-binder | — |
| 143 | MIMOSA<sub>BG</sub> | Sequence-fixed | 8IC0 | 8coh | Yes | 8 | 8 | 24 | Non-binder | — |
| 144 | MIMOSA<sub>BG</sub> | Sequence-fixed | 8IC0 | 8z8v | Yes | 10 | 9 | 6 | Non-binder | — |
| 145 | MIMOSA<sub>BG</sub> | Sequence-fixed | 8IC0 | 8coh | Yes | 10 | 6 | 24 | Non-binder | — |
| 146 | MIMOSA<sub>BG</sub> | Sequence-fixed | 8IC0 | 8coh | Yes | 7 | 7 | 20 | Non-binder | — |
| 147 | MIMOSA<sub>BG</sub> | Sequence-fixed | 8IC0 | 8coh | Yes | 7 | 6 | 20 | Non-binder | — |
| 148 | MIMOSA<sub>BG</sub> | Sequence-fixed | 8IC0 | 8coh | Yes | 9 | 9 | 21 | Non-binder | — |
| 149 | MIMOSA<sub>BG</sub> | Sequence-fixed | 8IC0 | 3eak | Yes | 11 | 8 | 15 | Non-binder | — |
| 150 | MIMOSA<sub>BG</sub> | Sequence-fixed | 8IC0 | 3eak | Yes | 11 | 8 | 17 | Non-binder | — |
| 151 | MIMOSA<sub>BG</sub> | Sequence-fixed | 8IC0 | 8coh | Yes | 8 | 10 | 21 | Non-binder | — |
| 152 | MIMOSA<sub>BG</sub> | Sequence-fixed | 8IC0 | 3eak | Yes | 10 | 8 | 13 | Non-binder | — |
| 153 | MIMOSA<sub>BG</sub> | Sequence-fixed | 8IC0 | 8coh | Yes | 7 | 10 | 21 | Non-binder | — |
| 154 | MIMOSA<sub>BG</sub> | Sequence-fixed | 8IC0 | 8coh | Yes | 9 | 6 | 21 | Non-binder | — |
| 155 | MIMOSA<sub>BG</sub> | Sequence-fixed | 8IC0 | 3eak | Yes | 9 | 8 | 14 | Non-binder | — |
| 156 | MIMOSA<sub>BG</sub> | Sequence-fixed | 8IC0 | 8coh | Yes | 8 | 8 | 21 | Non-binder | — |
| 157 | MIMOSA<sub>BG</sub> | Sequence-fixed | 8IC0 | 8coh | Yes | 8 | 8 | 21 | Non-binder | — |
| 158 | MIMOSA<sub>BG</sub> | Sequence-fixed | 8IC0 | 8z8v | Yes | 10 | 9 | 6 | Non-binder | — |
| 159 | MIMOSA<sub>BG</sub> | Sequence-fixed | 8IC0 | 8coh | Yes | 7 | 9 | 21 | Non-binder | — |
| 160 | MIMOSA<sub>BG</sub> | Sequence-fixed | 8IC0 | 8coh | Yes | 7 | 9 | 21 | Non-binder | — |
| 161 | MIMOSA<sub>BG</sub> | Sequence-fixed | 8IC0 | 8coh | Yes | 8 | 8 | 15 | Non-binder | — |
| 162 | MIMOSA<sub>BG</sub> | Sequence-fixed | 8IC0 | 8coh | Yes | 7 | 8 | 22 | Non-binder | — |
| 163 | MIMOSA<sub>BG</sub> | Sequence-fixed | 8IC0 | 8coh | Yes | 9 | 9 | 15 | Non-binder | — |
| 164 | MIMOSA<sub>BG</sub> | Sequence-fixed | 8IC0 | 8coh | Yes | 9 | 7 | 23 | Non-binder | — |
| 165 | MIMOSA<sub>BG</sub> | Sequence-fixed | 8IC0 | 8coh | Yes | 7 | 6 | 20 | Non-binder | — |
| 166 | MIMOSA<sub>BG</sub> | Sequence-fixed | 8IC0 | 8coh | Yes | 6 | 9 | 21 | Non-binder | — |
| 167 | MIMOSA<sub>BG</sub> | Sequence-fixed | 8IC0 | 3eak | Yes | 10 | 9 | 15 | Non-binder | — |
| 168 | MIMOSA<sub>BG</sub> | Sequence-fixed | 8IC0 | 8coh | Yes | 7 | 8 | 20 | Non-binder | — |
| 169 | MIMOSA<sub>BG</sub> | Sequence-fixed | 8IC0 | 8coh | Yes | 7 | 8 | 20 | Non-binder | — |
| 170 | MIMOSA<sub>BG</sub> | Sequence-fixed | 8IC0 | 3eak | Yes | 11 | 8 | 15 | Non-binder | — |
| 171 | MIMOSA<sub>BG</sub> | Sequence-fixed | 8IC0 | 8z8v | Yes | 7 | 7 | 10 | Non-binder | — |
| 172 | MIMOSA<sub>BG</sub> | Sequence-fixed | 8IC0 | 8z8v | Yes | 10 | 8 | 6 | Non-binder | — |
| 173 | MIMOSA<sub>BG</sub> | Sequence-fixed | 8IC0 | 3eak | Yes | 11 | 10 | 13 | Non-binder | — |
| 174 | MIMOSA<sub>BG</sub> | Sequence-fixed | 8IC0 | 8coh | Yes | 7 | 9 | 21 | Non-binder | — |
| 175 | MIMOSA<sub>BG</sub> | Sequence-fixed | 8IC0 | 3eak | Yes | 9 | 8 | 20 | Non-binder | — |
| 176 | MIMOSA<sub>BG</sub> | Sequence-fixed | 8IC0 | 8coh | Yes | 9 | 8 | 21 | Non-binder | — |
| 177 | MIMOSA<sub>BG</sub> | Sequence-fixed | 8IC0 | 8coh | Yes | 9 | 6 | 18 | Non-binder | — |
| 178 | MIMOSA<sub>BG</sub> | Sequence-fixed | 8IC0 | 8coh | Yes | 6 | 6 | 20 | Non-binder | — |
| 179 | MIMOSA<sub>BG</sub> | Sequence-fixed | 8IC0 | 3eak | Yes | 9 | 7 | 20 | Not assayable | — |
| 180 | MIMOSA<sub>BG</sub> | Sequence-fixed | 8IC0 | 3eak | Yes | 11 | 8 | 15 | Not assayable | — |

### B10. Computational overhead of mimic-guided sampling

**Backbone generation.** We measured backbone-generation time in a controlled $2 \times 2$ ablation of unguided sampling, Feynman–Kac (FK) steering, backward gradient guidance, and their combination. For each target and condition, ten backbones were generated as one batch on a single NVIDIA B200 GPU, using 500 reverse steps and four FK particles. The measurements cover the prediction loop only.

Table S15: Backbone-generation time for BoltzGen and MIMOSA<sub>BG</sub> component ablations on one B200 (seconds per design). Fold overhead relative to matched unguided sampling is shown in parentheses; each cell is one ten-design batch.

| Target | Unguided | FK only | Gradient only | FK + gradient |
| --- | --- | --- | --- | --- |
| KEAP1 | 2.70 (1.00×) | 6.80 (2.52×) | 3.30 (1.22×) | 7.80 (2.89×) |
| uPA | 2.50 (1.00×) | 6.20 (2.48×) | 3.10 (1.24×) | 7.10 (2.84×) |
| IL-8 | 1.90 (1.00×) | 3.60 (1.89×) | 2.30 (1.21×) | 4.30 (2.26×) |

Backward gradient guidance added 21–24% to unguided runtime, FK steering increased runtime to 1.89–2.52×, and combining both components resulted in a 2.26–2.89× overhead. The four FK particles ran in parallel on the GPU, while reverse steps ran sequentially. Backbone generation for a full 60,000-design KEAP1 MIMOSA<sub>BG</sub> cohort would cost 130.0 B200-hours, versus 45.0 unguided. For a full 100,000-design IL-8 MIMOSA<sub>BG</sub> cohort, it would cost 119.4 versus 52.8 unguided.

**Including inverse folding and refolding.** Backbone generation, inverse folding, and refolding together produce an evaluable design candidate, making the full pipeline the relevant unit for measuring guidance overhead. For each of the 120 ablation backbones, we generated one inverse-folded sequence and refolded it with 200 diffusion steps and five structure samples. Because these downstream stages were configured identically across guidance conditions, their times were pooled within each target and added to the corresponding backbone-generation times (Table S16). This shared downstream cost reduces the relative overhead of guidance but not its absolute cost. The estimates exclude filtering, retries, failed-design replacement, independent structure-prediction oracles, and analysis.

Table S16: Full-pipeline time (generation, inverse folding, and refolding) for BoltzGen and MIMOSA<sub>BG</sub> component ablations on one B200 (seconds per design). The common downstream cost was pooled across guidance conditions within each target. Fold overhead relative to the matched unguided pipeline is shown in parentheses.

| Target | Unguided | FK only | Gradient only | FK + gradient |
| --- | --- | --- | --- | --- |
| KEAP1 | 15.03 (1.00×) | 19.13 (1.27×) | 15.63 (1.04×) | 20.13 (1.34×) |
| uPA | 14.08 (1.00×) | 17.78 (1.26×) | 14.68 (1.04×) | 18.68 (1.33×) |
| IL-8 | 9.55 (1.00×) | 11.25 (1.18×) | 9.95 (1.04×) | 11.95 (1.25×) |
